# Purinergic signaling in cochlear supporting cells reduces hair cell excitability by increasing the extracellular space

**DOI:** 10.1101/852756

**Authors:** Travis A. Babola, Calvin J. Kersbergen, Han Chin Wang, Dwight E. Bergles

**Affiliations:** The Solomon Snyder Department of Neuroscience, Johns Hopkins University, Baltimore, Maryland 21205, USA; Department of Otolaryngology Head and Neck Surgery, Johns Hopkins University, Baltimore, Maryland 21287, USA; Johns Hopkins University Kavli Neuroscience Discovery Institute, Baltimore, Maryland, 21205

## Abstract

Neurons in developing sensory pathways exhibit spontaneous bursts of electrical activity that are critical for survival, maturation and circuit refinement. In the auditory system, intrinsically generated activity arises within the cochlea, but the molecular mechanisms that initiate this activity remain poorly understood. We show that burst firing of mouse inner hair cells prior to hearing onset requires P2RY1 autoreceptors expressed by inner supporting cells. P2RY1 activation triggers K^+^ efflux and depolarization of hair cells, as well as osmotic shrinkage of supporting cells that dramatically increased the extracellular space and speed of K^+^ redistribution. Pharmacological inhibition or genetic disruption of P2RY1 suppressed neuronal burst firing by reducing K^+^ release, but unexpectedly enhanced their tonic firing, as water resorption by supporting cells reduced the extracellular space, slowing K^+^ clearance. These studies indicate that purinergic signaling in supporting cells regulates hair cell excitability by controlling the volume of the extracellular space.

## Introduction

The developing nervous system must generate, organize, and refine billions of neurons and their connections. While molecular guidance cues forge globally precise neuronal connections between distant brain areas (***Stoeckli, 2018***; ***Dickson, 2002***), the organization of local connections is initially coarse and imprecise (***Dhande et al., 2011***; ***Kirkby et al., 2013***; ***Sretavan and Shatz, 1986***). Coincident with the refinement of topographic maps, nascent circuits experience bursts of intrinsically generated activity that emerge before sensory systems are fully functional (***Kirkby et al., 2013***). This intrinsically generated activity consists of periodic bursts of high frequency firing that promotes the survival and maturation of neurons in sensory pathways (***Blankenship and Feller, 2010***; ***Moody and Bosma, 2005***). The precise patterning of this electrical activity appears crucial for refinement of local connections, as its disruption results in improper formation of topographic maps (***Antón-Bolaños et al., 2019***; ***Burbridge et al., 2014***; ***Xu et al., 2011***) and impaired maturation and specification of sensory neurons (***Shrestha et al., 2018***; ***Sun et al., 2018***). In all sensory systems that have been examined, spontaneous burst firing arises within their respective developing sensory organs, e.g. retina, olfactory bulb, spindle organ, and cochlea (***Blankenship and Feller, 2010***). Although the mechanisms that induce spontaneous activity in the developing retina have been ex-tensively explored, much less is known about the key steps involved in triggering auditory neuron burst firing in the developing cochlea. Understanding these processes may provide novel insights into the causes of developmental auditory disorders, such as hypersensitivity to sounds and auditory processing disorders that prevent children from communicating and learning effectively.

The mechanisms responsible for initiating spontaneous activity appear to be unique to each sensory system, reflecting adaptations to the structure and cellular composition of the sensory organs. In the cochlea, two distinct models have been proposed to initiate burst firing of inner hair cells. One model proposes that burst firing results from intermittent hyperpolarization of tonically active IHCs by cholinergic efferents (***Johnson et al., 2011***; ***Wang and Bergles, 2014***), which provide prominent inhibitory input to IHCs prior to hearing onset (***Glowatzki and Fuchs, 2000***). Consistent with this model, activation of acetylcholine receptors in acutely isolated cochleae caused IHCs to switch from sustained to burst firing (***Johnson et al., 2011***). However, *in vivo* recordings from auditory brainstem revealed that neuronal burst firing remains, with altered features, in α9 knockout mice (***Clause et al., 2014***), which lack functional efferent signaling in IHCs (***Johnson et al., 2013***), and persists in IHCs and auditory neurons in cochleae maintained *in vitro* without functional efferents (***Johnson et al., 2013***). Thus, cholinergic efferents appear to modulate the temporal characteristics of bursts, but are not essential to initiate each event.

An alternative model proposes that IHCs are induced to fire bursts of action potentials by the release of K^+^ from nearby inner supporting cells (ISCs), which together form a transient structure known as Köllikers organ (Greater Epithelial Ridge) that is prominent prior to hearing onset (***Tritsch et al., 2010b***). K^+^ release from ISCs occurs following a cascade of events that begins with the spontaneous release of ATP and activation of purinergic autoreceptors. Purinergic receptor activation induces an increase in intracellular Ca^2+^, opening of Ca^2+^-activated Cl^−^ channels (TMEM16A), and efflux of Cl^−^ and subsequently K^+^ to balance charge (***Tritsch et al., 2007***; ***Wang et al., 2015***). The loss of ions during each event draws water out of ISCs through osmosis, leading to pronounced shrinkage (crenation) of ISCs. While these pathways have been extensively studied *in vitro*, the molecular identity of the purinergic receptors has remained elusive and few manipulations of key steps in this pathway have been performed *in vivo*, limiting our understanding of how spontaneous activity is generated at this critical stage of development.

Here, we show that the key initial step in generation of spontaneous activity in the auditory system involves activation of P2RY1 autoreceptors in ISCs. These metabotropic receptors induce Ca^2+^ release from intracellular stores that allow TMEM16A channels to open. Pharmacological inhibition of P2RY1 or genetic deletion of *P2ry1* dramatically reduced burst firing in spiral ganglion neurons (SGNs) and blocked the coordinated, spatially restricted activation of ISCs, IHCs, and SGNs in the cochlea. Unexpectedly, P2RY1 activation also promoted the clearance of K^+^ by increasing the volume of extracellular space, enhancing the diffusion of K^+^ ions away from IHCs. Conversely, inhibition of P2RY1 reduced the extracellular space and restricted the redistribution of K^+^ within the cochlear epithelium, causing IHCs to depolarize and fire tonically, demonstrating an important role for purinergic receptor-mediated extracellular space changes in controlling IHC excitability. Using *in vivo* widefield epifluorescence imaging of the auditory midbrain in unanesthetized mice, we show that acute inhibition of P2Y1 receptors dramatically reduced burst firing of auditory neurons in isofrequency domains. Together, these data indicate P2RY1 autoreceptors in non-sensory, cochlear supporting cells play a crucial role in generating bursts of activity among neurons that will ultimately process similar frequencies of sound, providing the means to initiate the maturation of auditory pathways before hearing onset.

## Results

### Supporting cell spontaneous currents require calcium release from intracellular stores

Periodic release of ATP from ISCs in the developing cochlea initiates a signaling cascade in these cells that increases intracellular calcium (Ca^2+^), opens Ca^2+^-activated Cl^−^ channels (TMEM16A), and ultimately results in efflux of chloride and K^+^ into the extracellular space. Although the increase in intracellular Ca^2+^ following activation of purinergic autoreceptors is sufficient to induce both depolarization and osmotic shrinkage (crenation; ***Wang et al.*** (***2015***)), the relative contributions of Ca^2+^ influx (e.g. through Ca^2+^-permeable, ionotropic P2X receptors) and release from intracellular stores (e.g. following metabotropic P2Y receptor activation) to these cytosolic Ca^2+^ transients is unclear. To define the signaling pathways engaged by purinergic receptor activation, we examined the sensitivity of spontaneous ISC whole-cell currents and crenations to inhibitors of intracellular Ca^2+^ pathways (***Figure 1***A). Spontaneous inward currents and crenations were abolished following a 15 minute incubation of excised cochlea in BAPTA-AM (100µM), a cell permeant Ca^2+^ chelator (***Figure 1***B-F), and after depleting intracellular Ca^2+^ stores with thapsigargin (2µM), an inhibitor of endoplasmic reticulum Ca^2+^-ATPase (***Figure 1***B-F). These data suggest that Ca^2+^ release from intra-cellular stores is necessary for spontaneous electrical activity in ISCs.

**Figure 1.**
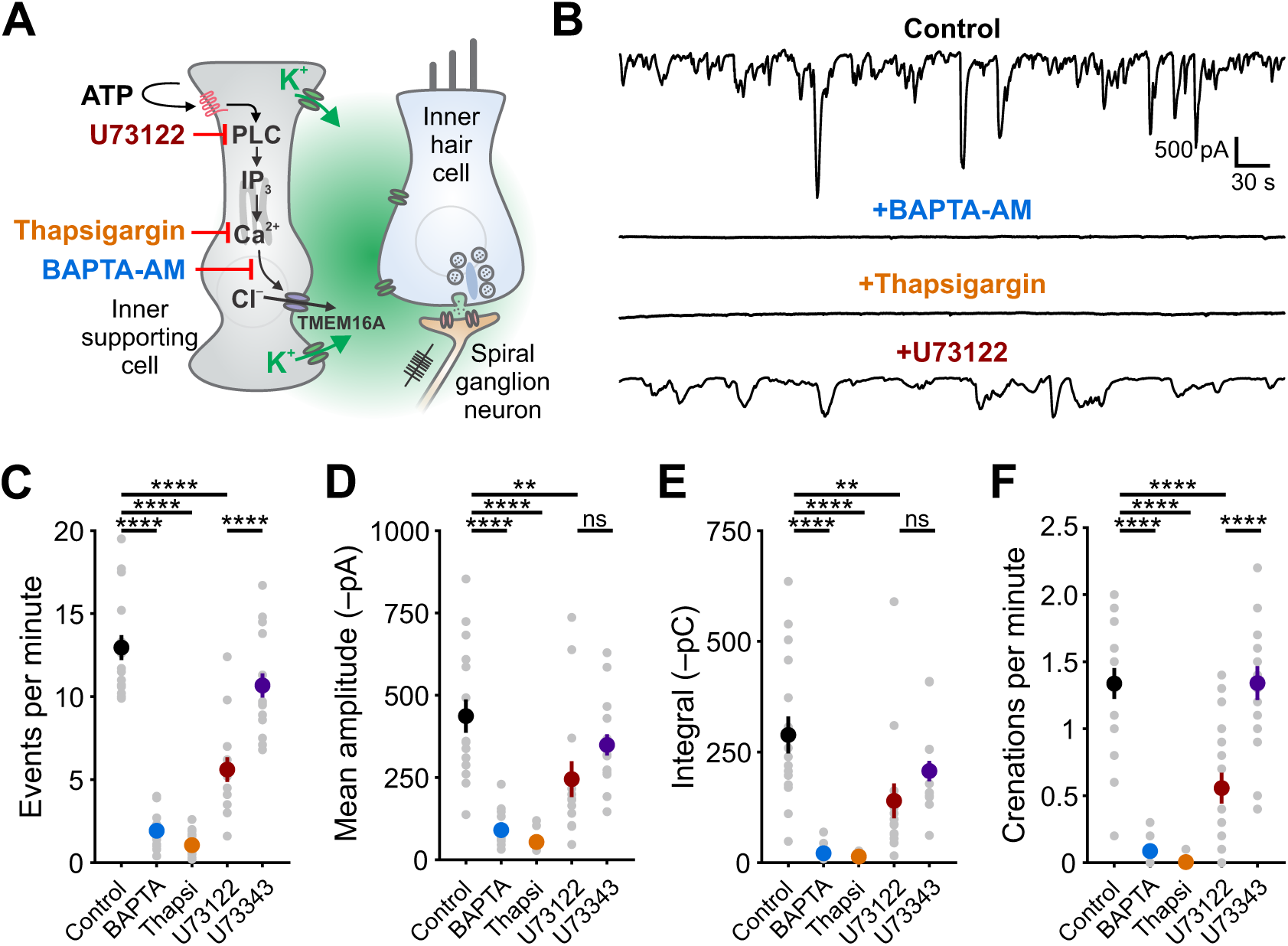
Ca^2+^ release from intracellular stores is required for spontaneous currents and crenation in inner supporting cells. **(A)** Model of ATP-mediated depolarization of inner hair cells. ATP: adenosine triphosphate, PLC: phospholipase C, IP3: inositol triphosphate, TMEM16A: transmembrane member 16A (Ca^2+^-activated Cl^−^channel). Inhibitors of key steps in this pathway are indicated. **(B)** Whole-cell voltage-clamp recordings from inner supporting cells after pre-incubating with indicated inhibitors. **(C)** Quantification of ISC spontaneous current frequency in the presence of inhibitors of the G_q_ pathway. Data shown as mean ± SEM. n = 16 cells (control), 16 cells (BAPTA-AM; 100µM), 20 cells (Thapsigargin; 2µM), 14 cells (U73211; 10µM), and 16 cells (U73343; 10µM). ****p<5e-5, one-way ANOVA. **(D)** Quantification of ISC spontaneous current amplitude in the presence of inhibitors of the G_q_ pathway. Data shown as mean ± SEM. n values are reported in (C) (one-way ANOVA; ****p<5e-5, **p<0.005, ns: not significant). **(E)** Quantification of ISC spontaneous current charge transfer (integral) in the presence of inhibitors of the G_q_ pathway. Data shown as mean ± SEM. N values are reported in (C) (one-way ANOVA; ****p<5e-5, **p<0.005, ns: not significant). **(F)** Quantification of ISC crenation (cell shrinkage) frequency in the presence of inhibitors of the G_q_ pathway. Data shown as mean ±SEM. n values are reported in (C) (one-way ANOVA; ****p<5e-5, **p<0.005, ns: not significant).

Metabotropic G_q_-coupled receptors typically induce PLC-mediated cleavage of phosphatidyli-nositol 4,5-bisphosphate (PIP_2_) and subsequent binding of inositol trisphosphate (IP_3_) to IP_3_ receptor-channels on the endoplasmic reticulum to release Ca^2+^ into the cytoplasm. To investigate if PLC signaling is required for generation of spontaneous activity in ISCs, we recorded spontaneous currents and crenations from ISCs in the presence of U73122 (10µM), a PLC inhibitor, and U73343 (10µM), an inactive succinimide analog. The frequency of spontaneous currents and crenations were dramatically reduced following U73122 incubation, but not U73343 (***Figure 1***B-F); the amplitudes and charge transfer of residual activity also trended lower during PLC inhibition, but this did not reach significance due to high variance in the sizes of these responses (***Figure 1***B-F). Together, these results suggest that engagement of a G_q_-coupled purinergic autoreceptor is a critical first step in initiating PLC-mediated Ca^2+^ release from intracellular stores and subsequent activation of TMEM16A channels.

### The metabotropic purinergic receptor P2Y1 is highly expressed by supporting cells

There are eight members of the metabotropic purinergic receptor family in mouse, four of which are G_q_-coupled (P2RY1, P2RY2, P2RY4, and P2RY6). Gene expression studies in the developing mouse cochlea revealed that non-sensory cells express *P2ry1* mRNA at high levels (***Scheffer et al., 2015***), >100 fold higher than any other P2ry (***Figure 2***A) and that expression of this receptor progressively increases during early postnatal development (***Figure 2***A, inset) concurrent with increases in spontaneous activity (***Tritsch and Bergles, 2010***). To determine which cells in the sensory epithelium express P2RY1, we isolated cochleae from *P2ry1-LacZ* reporter mice and performed X-gal staining. Intense blue labeling was present along the entire length of the cochlea within Kölliker’s organ (Greater Epithelial Ridge; ***Figure 2***B). Cross-sections of cochlea revealed that staining was present within ISCs, but not IHCs (Myosin VIIA, ***Figure 2***C), indicating that P2RY1 is properly localized to sense ATP release from ISCs prior to hearing onset.

**Figure 2.**
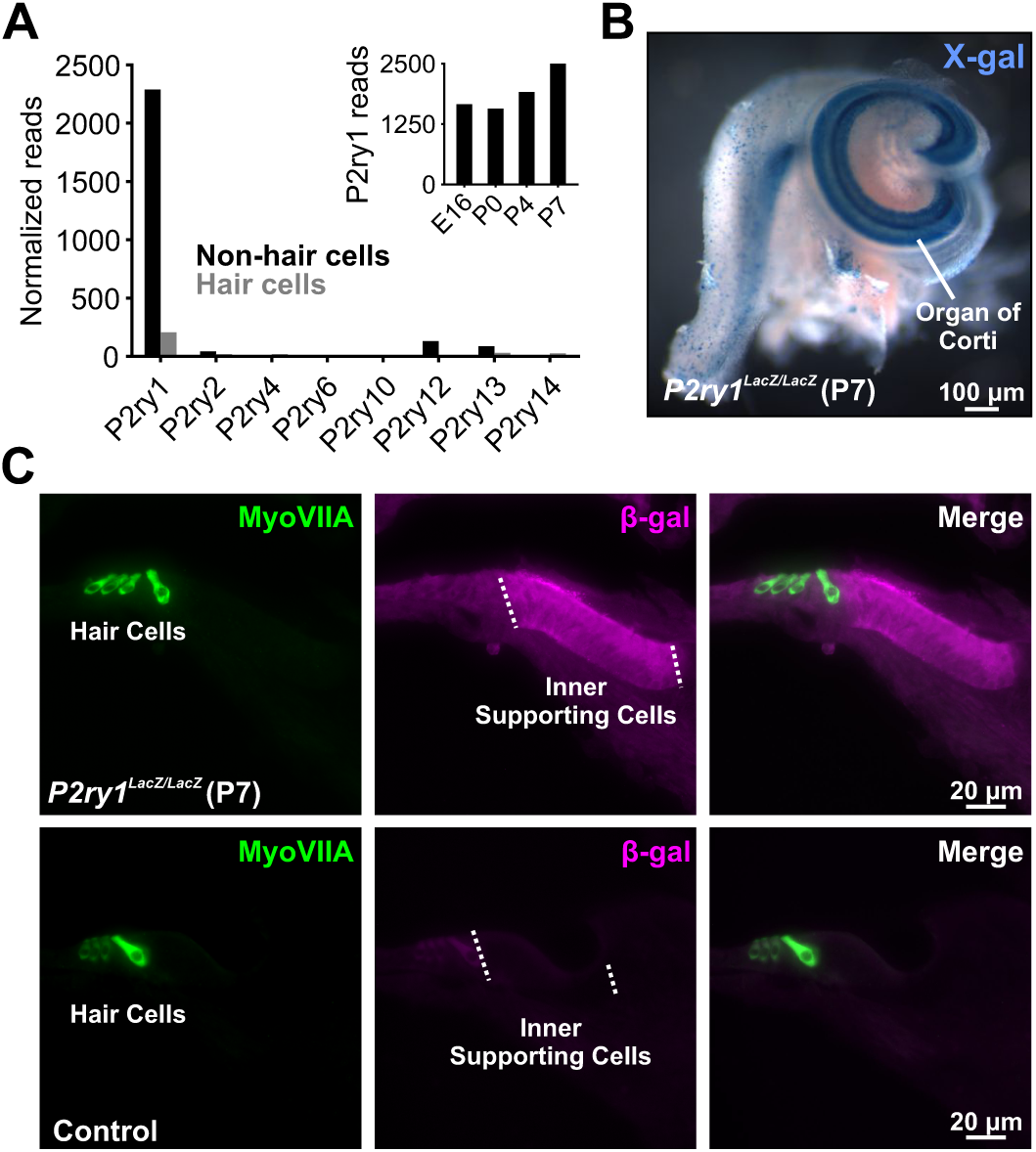
The metabotropic P2Y1 receptor is highly expressed by ISCs. **(A)** Expression levels of metabotropic purinergic receptors in hair cells (grey) and non-sensory cells (black) of the developing cochlea (postnatal day 7, P7). (inset) *P2ry1* expression in non-sensory cells over development. Data adapted from ***Scheffer et al.*** (***2015***). **(B)** Image of a cochlea following X-gal reaction in *P2ry1-LacZ* reporter mice. **(C)** Immunostaining for β-galactosidase in cochleae from P7 *P2ry1-LacZ* (top) and control (bottom) cochlea.

### P2RY1 signaling is required for spontaneous activity in ISCs and IHCs

To determine if P2RY1 is responsible for spontaneous ATP-mediated currents in ISCs, we examined the sensitivity of these responses and associated crenations to the P2RY1 antagonist MRS2500 (***Figure 3***A,B). Acute inhibition of P2RY1 with MRS2500 (1µM) markedly reduced both spontaneous ISC currents (***Figure 3***B,C) and crenations (***Figure 3***D,E); near complete inhibition occurred within minutes at both room temperature (***Figure 3***B,C) and near physiological temperature (***Figure 3–Figure Supplement 1***A-G), with only sporadic, small amplitude events remaining that were not mediated by purinergic receptors (***Figure 3–Figure Supplement 1***B-E). Consistent with the involvement of P2RY1, the amplitude and total charge transfer of ISC events (***Figure 3–Figure Supplement 2***A,B) and size of spontaneous crenations (***Figure 3–Figure Supplement 2***C,D) were smaller in cochleae in *P2ry1* KO mice relative to controls. However, supporting cells *P2ry1* KO mice exhibited aberrant, gain-of-function activity consisting of frequent, small amplitude currents (***Figure 3–Figure Supplement 2***A,B), that were not blocked by MRS2500 or broad-spectrum P2 receptor antagonists (***Figure 3–Figure Supplement 2***E,F).

**Figure 3.**
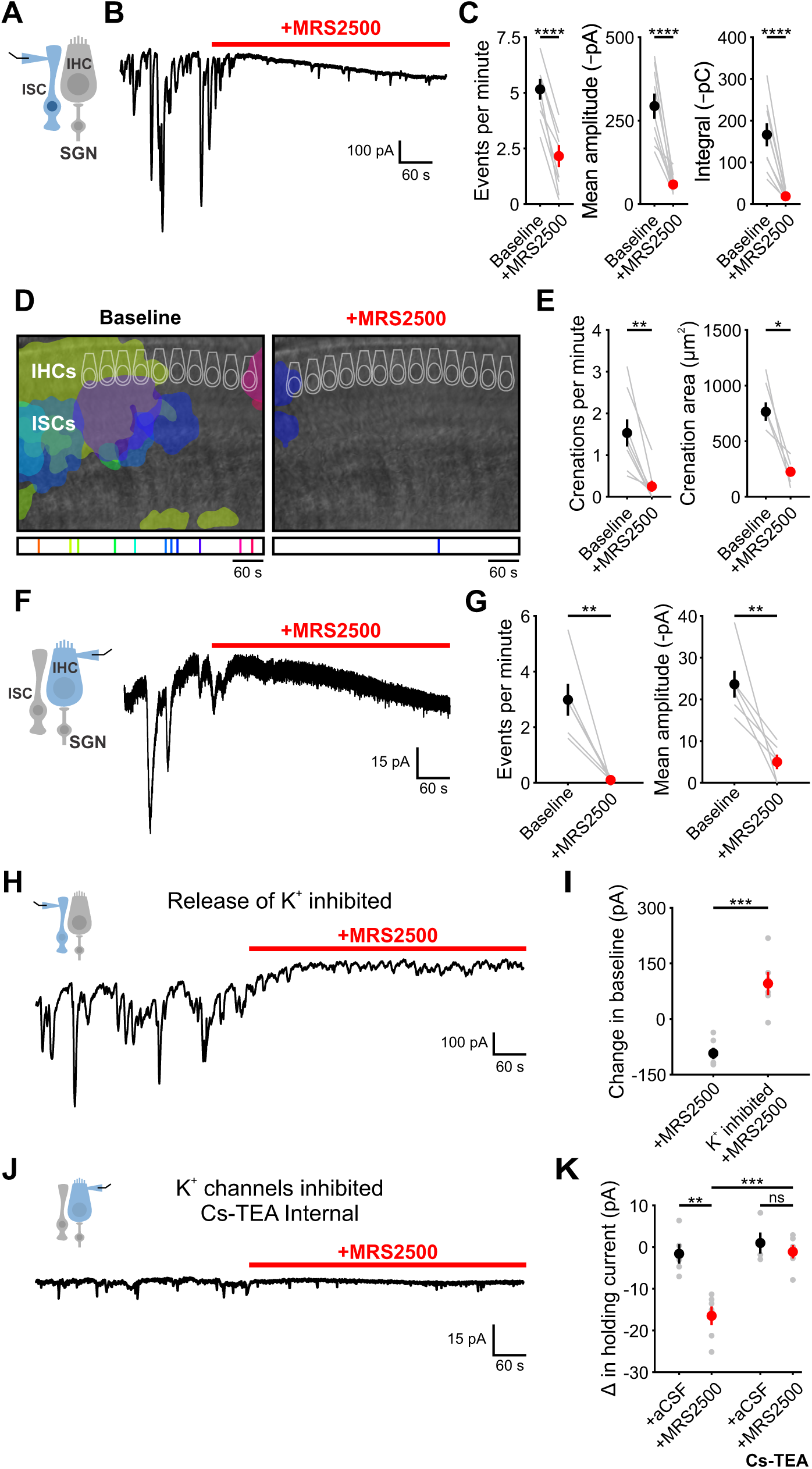
P2Y1 inhibition abolishes spontaneous currents in inner supporting cells and inner hair cells. **(A)** Schematic of ISC whole-cell recording configuration. **(B)** ISC spontaneous inward currents before and after application of MRS2500 (1µM). Recordings performed at ∼25°C. **(C)** Plot of event frequency, amplitude, and integral (charge transfer) before and after MRS2500 application. n = 8 ISCs (two-tailed paired Student’s t test; ****p<5e-5) **(D)** Intrinsic optical imaging performed before and after application of the P2RY1 antagonist, MRS2500 (1µM). Detected crenations are outlined in colors based on time of occurrence as indicated below image. Imaging performed at ∼25°C. Caption continued on next page.

ATP-mediated signaling in ISCs activates TMEM16A, triggering K^+^ efflux that depolarizes nearby IHCs. To assess whether P2RY1 signaling is also required for periodic excitation of IHCs prior to hearing onset, we assessed the sensitivity of spontaneous IHC inward currents to MRS2500 (***Figure 3***F). Consistent with the supporting cell origin of IHC activity, application of MRS2500 (1µM) also abolished spontaneous currents in IHCs (***Figure 3***F,G). Together, these data suggest that P2RY1 is the primary purinergic autoreceptor on ISCs responsible for inducing periodic excitation of hair cells prior to hearing onset.

### P2RY1 inhibition leads to extracellular K^+^ accumulation

Although P2RY1 inhibition abolished most transient inward currents in both ISCs and IHCs, a progressively increasing inward current (downward shift in baseline) appeared in both cell types with prolonged application of MRS2500 (***Figure 3***B,F). Prior studies in CNS brain slices indicated that G_q_-coupled purinergic receptors in astrocytes regulate extracellular K^+^ concentration and neuronal excitability (***Wang et al., 2012***). The slowly progressing nature of the response in IHCs and ISCs suggest that it may similarly arise from accumulation of K^+^ released from cells in the organ of Corti. If this hypothesis is correct, then inhibiting the main sources of K^+^ should diminish this inward current. Indeed, when IHC and SGN excitation was inhibited with tetrodotoxin (TTX, 1µM) and cadmium (CdCl_2_, 100µM), and the K^+^ transporters, Na,K-ATPase and NKCC, were inhibited with ouabain (10µM) and bumetanide (50µM), no inward current was induced in ISCs upon blocking P2RY1 (***Figure 3***H,I). Similarly, if K^+^ accumulation is responsible for the current in IHCs, it should be abolished when the ability of IHCs to detect changes in K^+^ is reduced. When whole cell recordings were performed from IHCs using an internal solution containing Cs^+^ and TEA, which blocks most IHC K^+^ channels (***Kros et al., 1998***; ***Marcotti et al., 2003***), MRS2500 also did not induce an inward current (***Figure 3***J,K). Together, these results suggest that P2RY1 has two distinct effects in the cochlea; it induces the transient inward currents that triggers IHC burst firing and it accelerates the clearance of K^+^ within the organ of Corti.

To directly assess the relationship between P2RY1 activity and extracellular K^+^ accumulation near IHCs, we monitored K^+^ levels in the extracellular space using IHC K^+^ channels. Focal P2RY1 stimulation with a selective agonist (MRS2365, 10µM), which mimics the effect of endogenous ATP by eliciting an inward current and crenations in ISCs in control but not *P2ry1* KO mice (***Figure 4***A-C), was combined with assessments of the reversal potential of K^+^ currents in IHCs using a voltage protocol similar to one used to assess extracellular K^+^ buildup at vestibular calyceal synapses (***Lim et al., 2011***). This protocol consisted of: (1) a hyperpolarizing step to –110mV to relieve K^+^ channel inactivation, (2) a depolarizing step to +30mV to activate outward K^+^ currents, and (3) a step to –70mV to obtain a “tail” current (***Figure 4***D-F). Because the conductance during this last step is largely mediated by K^+^ channels, it is highly sensitive to shifts in K^+^ driving force induced by changes in extracellular K^+^ (***Contini et al., 2017***; ***Lim et al., 2011***). Following transient stimulation of P2RY1, K^+^ tail currents immediately shifted inward, as would be expected if extracellular K^+^ increases (***Figure 4***G,I), and is similar to the effects of a metabotropic purinergic receptor agonist (UTP) on synaptically-evoked K^+^ currents in IHCs (***Wang et al., 2015***). However, after a few seconds these K^+^ currents shifted outward relative to baseline, indicative of a gradual decrease in extracellular K^+^ below that present prior to P2RY1 stimulation, then gradually returned to the pre-stimulation level after several minutes (***Figure 4***G,I).

**Figure 4.**
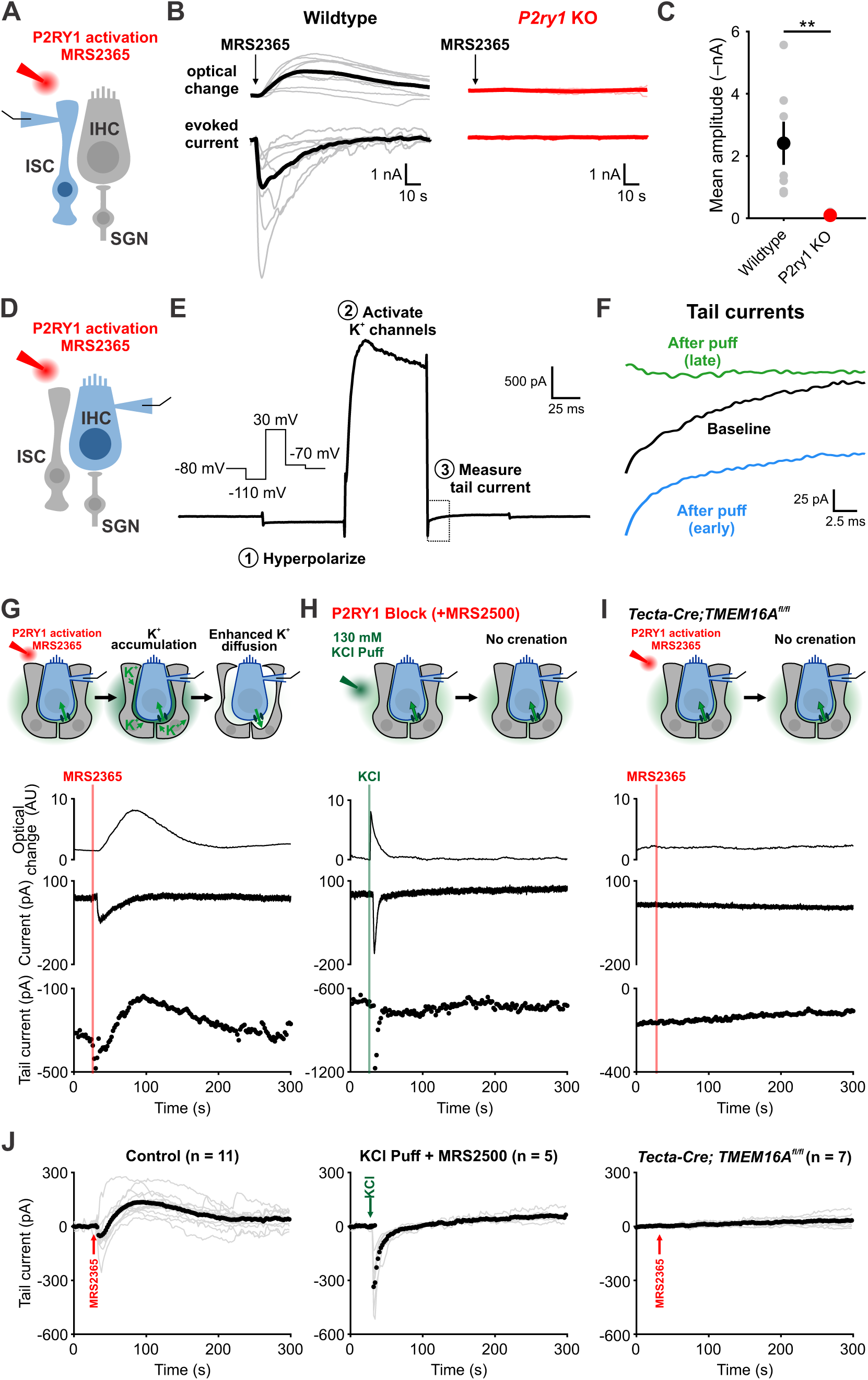
Caption on next page. **Figure 4.** Activation of P2RY1 results in an initial accumulation of extracellular K^+^, followed by crenation and enhanced K^+^ clearance. **(A)** Schematic of whole-cell recording configuration from ISCs with puffs of MRS2365 (10µM), a P2RY1 agonist. **(B)** Optical change (crenation) and current elicited with MRS2365 puffs in wildtype and *P2ry1* KO mice. **(C)** Plot of mean current amplitude with MRS2365 puffs. n = 6 and n = 5 ISCs from wildtype and *P2ry1* KO mice, respectively (two-tailed Student’s t test; **p<0.005). **(D)** Schematic of whole-cell recording configuration from IHCs with puffs of MRS2365 (10µM). **(E)** Example current trace and voltage-protocol designed to measure K^+^ accumulation. Dashed box indicated tail current measurement period indicated in (F). **(F)** Tail currents observed during baseline, immediately following the MRS2365 puff (with 2 seconds), and after the puff (30 seconds). **(G)** Model of K^+^ dynamics following MRS2365 stimulation. Initially, extracellular K^+^ rapidly increases following stimulation, but ISCs crenate, increasing the amount of extracellular space and K^+^ buffering. (bottom) Optical change (crenation), holding current, and tail current as a function of time with respect to MRS2365 puff. **(H)** Similar to G, but with KCl puffs (130µM) in cochleae treated with MRS2500. **(I)** Similar to G, but in *Tecta-Cre;TMEM16A*^*fl/fl*^ mice where TMEM16A has been conditionally removed from the sensory epithelium (see ***Figure 4–Figure Supplement 1***). No crenations were observed with MRS2365 stimulation. **(J)** Plot of tail currents over time following MRS2365 stimulation. Grey lines indicate individual IHCs; black points indicate the mean across the population. Baseline was normalized to 0 pA for all traces.

The outward shift in K^+^ tail current followed the time course of the crenation (*τ*_decay_ = 100 ± 14s for tail currents and *τ*_decay_ = 38 ± 4s for crenations), suggesting that the shrinkage of cells induced by P2RY1 activation results in a prolonged increase in extracellular space that may allow greater dilution and more rapid redistribution of K^+^ in the organ of Corti. Alternatively, buildup of extracellular K^+^ alone may stimulate greater uptake. To determine if rapid increases in extracellular K^+^ or Cl^−^ were sufficient to stimulate K^+^ redistribution in the absence of crenation, we puffed KCl (130mM) into the supporting cell syncytium in the presence of P2RY1 antagonists (***Figure 4***H). As expected, this transient increase in extracellular K^+^ induced an inward shift in K^+^ tail currents and a brief optical change induced by fluid delivery; however, K^+^ tail currents rapidly returned to baseline and did not shift outward, suggesting that K^+^ (and Cl^−^) efflux are not sufficient to enhance K^+^ redistribution rates. In addition, we transiently stimulated P2RY1 in *Tecta-Cre;TMEM16A*^*fl/fl*^ mice, in which purinergic receptor activation is preserved, but crenations are abolished (***Wang et al., 2015***). In these mice, ISCs failed to crenate, IHCs did not depolarize, and K^+^ tail currents remained stable throughout the duration of the recording (***Figure 4***I). These results suggest that purinergic autoreceptors on ISCs influence extracellular K^+^ levels by both triggering K^+^ release and by altering K^+^ redistribution by controlling the size of the extracellular space.

### *P2ry1* mediates coordinated neuronal activation and precise burst firing of SGNs

To evaluate the role of P2RY1 in initiating coordinated cellular activity in the cochlea, we monitored large-scale activity patterns in excised cochleae from *Pax2-Cre;R26-lsl-GCaMP3* mice, which express GCaMP3 in nearly all cells of the inner ear. Time lapse imaging revealed that the spontaneous Ca^2+^ elevations that occur simultaneously within groups of ISCs, IHCs, and SGNs (***Tritsch and Bergles, 2010***; ***Zhang-Hooks et al., 2016***) were abolished following inhibition of P2RY1 with MRS2500 (***Figure 5***A-C) and were dramatically reduced in *P2ry1* KO mice (*Pax2-Cre;R26-lsl-GCaMP3;P2ry1*^*–/–*^) (***Figure 5–Figure Supplement 1***A,B). Moreover, in accordance with the progressive increase in extracellular K^+^ that follows P2RY1 inhibition, there was a gradual increase in spontaneous, uncoordinated Ca^2+^ transients in IHCs in the presence of MRS2500 (***Figure 5***D-F), suggesting that this K^+^ accumulation increases IHC firing. Similarly, IHCs in *P2ry1* KO mice displayed a higher level of uncorrelated hair cell Ca^2+^ transients (***Figure 5–Figure Supplement 1***C-E), indicative of enhanced excitability. Together, these results indicate that P2RY1 is required for coordinated activation of ISCs, IHCs, and SGNs before hearing onset and that P2RY1 inhibition leads to higher rates of uncorrelated activity.

**Figure 5.**
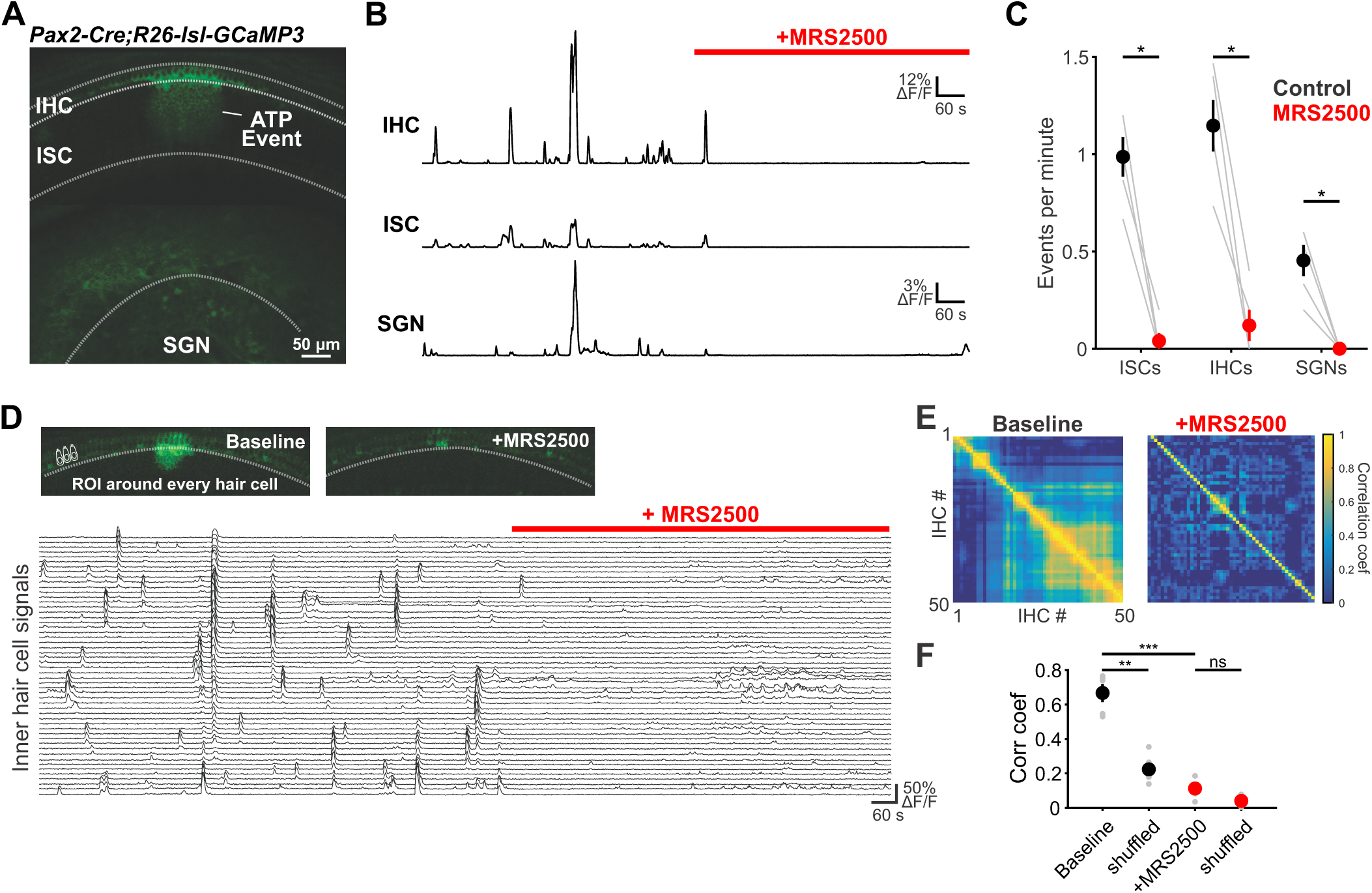
Large-scale coordinated activity in the cochlea requires P2RY1. **(A)** Exemplar Ca^2 +^ transient in excised cochlea from *Pax2-Cre;R26-lsl-GCaMP3* mice. Note coordinated activation of ISCs, IHCs, and SGNs. **(B)** Traces of fluorescence intensity over time taken from ROIs that span the entire IHC, ISC, and SGN regions indicated in (A). **(C)** Plot of event frequency before and during application of MRS2500 (1µM). n = 5 cochlea (two-tailed paired Student’s t test with Bonferroni correction; *p<0.05). **(D)** Exemplar images of IHC Ca^2+^ transients. ROIs were drawn around every IHC for subsequent analysis (bottom). **(E)** Correlation matrices generated by calculating the linear correlation coefficient for all IHC pairs before and after MRS2500 application. **(F)** Plot of average correlation coefficient calculated between the four nearest IHCs or four randomly shuffled IHCs. n = 5 cochleae (two-tailed paired Student’s t test with Bonferroni correction; ***p<0.0005, **p<0.005, ns, not significant).

IHCs in the developing cochlea exhibit regenerative Ca^2+^ spikes that strongly activate post-synaptic SGNs, resulting in bursts of action potentials that propagate to the CNS. To determine if P2RY1 initiates burst firing in SGNs, we recorded spontaneous activity from SGNs using juxta-cellular recordings from their somata (***Figure 6***A). Application of MRS2500 resulted in a dramatic reduction of high frequency burst firing in SGNs, visible as a decrease in burst frequency and action potentials per burst (***Figure 6***E,F). All SGN spiking was abolished by the AMPA receptor antagonist NBQX (50µM) (***Figure 6***D), indicating that their activity requires synaptic excitation by IHCs. The precise patterning of action potentials within bursts was also disrupted by P2RY1 inhibition, as there were fewer interspike intervals in the 75–125ms range (***Figure 6***C,F), which correspond to the maximum rate of Ca^2+^ spike generation by IHCs during ATP-mediated excitation (***Tritsch et al., 2010a***). Additionally, the coefficient of variation measured for interspike intervals was significantly lower following P2RY1 inhibition, suggesting SGNs fire more randomly (***Figure 6***E). However, the average frequency of action potentials remained unchanged during P2RY1 inhibition (***Figure 6***E), due to increases in non-burst firing (***Figure 3***F,***Figure 4***D). SGNs in *P2ry1* KO cochleae exhibited activity similar to wildtype SGNs in the presence of MRS2500, with a lower burst firing rate, fewer interspike intervals in the 75–125ms range, and a lower coefficient of variation of interspike intervals relative to controls (***Figure 6***G-I). However, despite the profound contribution of P2RY1 to ISC and IHC activity, some burst-like behavior was still observed in SGNs (***Figure 6***D,G), suggesting that other forms of excitation emerge in the absence of P2RY1, due to an increase in overall excitability or developmental changes. Together, these data indicate that P2RY1 is required to generate discrete bursts of action potentials in SGNs and that loss of these receptors enhances uncorrelated firing.

**Figure 6.**
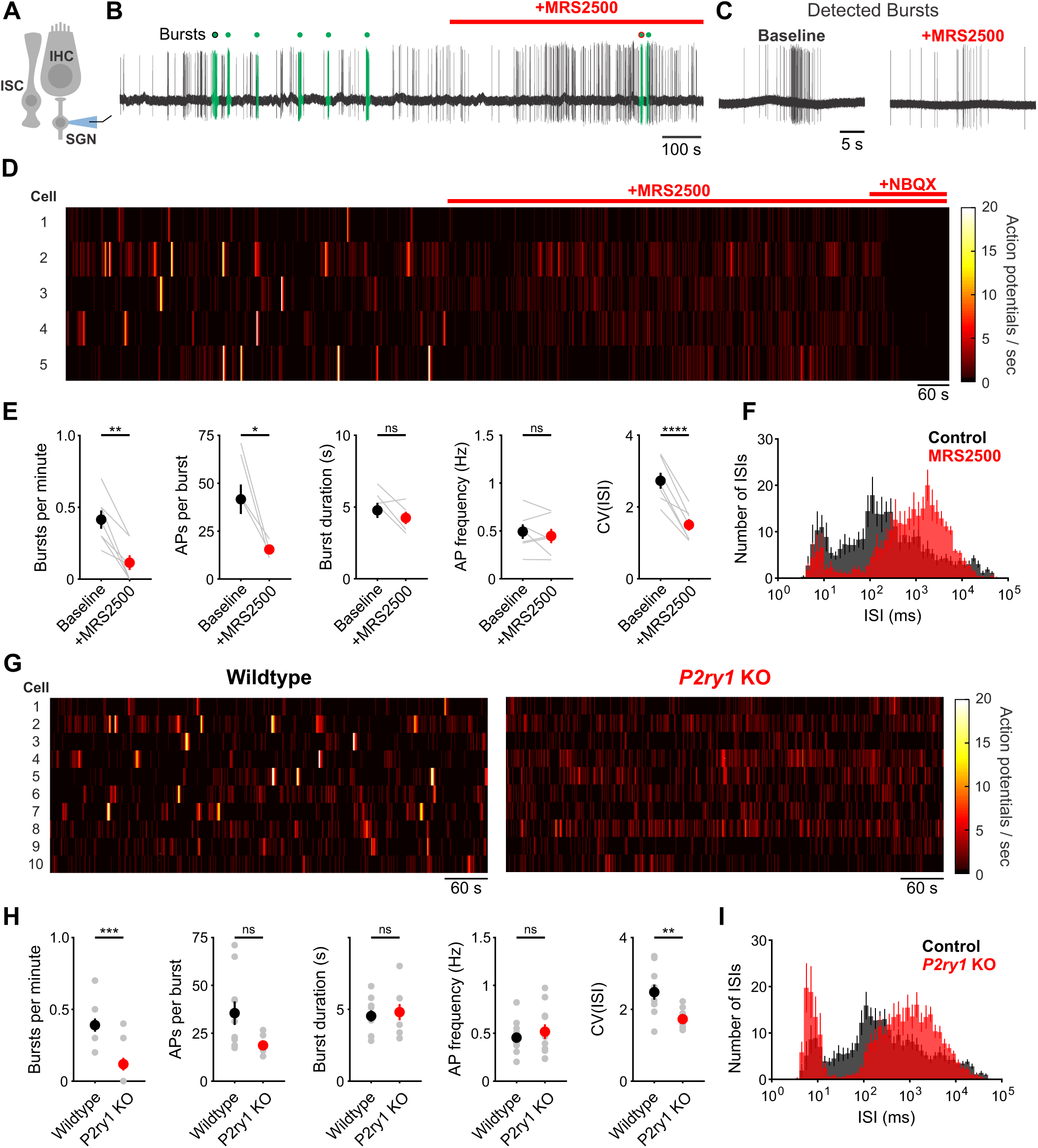
Inhibition of P2RY1 disrupts burst firing in spiral ganglion neurons. **(A)** Schematic of SGN juxtacellular recording configuration. All recordings were performed at ∼25°C. **(B)** Action potentials recorded before and during MRS2500 application (1µM). Detected bursts are indicated in green (see Methods and Materials for parameters used for burst detection). Circles with black and red outlines are expanded in (C).**(C)** Action potentials within a detected burst before and after MRS2500 application. **(D)** Raster plots indicating the average firing rate of SGNs (bin: 1 s) before and after MRS2500 application (1µM) and subsequent NBQX (50µM) **(E)** Plots of average burst frequency, burst duration, action potentials (AP) per burst, average AP frequency, and coefficient of variation for all interspike intervals (ISIs) measured. n = 7 SGNs from 7 cochleae (two-tailed paired Student’s t-test with Bonferroni correction; ****p<5e-5, **p<0.005, *p<0.05, ns, not significant). **(F)** Average log-binned interspike interval histograms before and after MRS2500 application. **(G)** Raster plots indicating the average firing rate of SGNs (bin: 1 s) in wildtype and *P2ry1* KO mice. **(H)** Plots of average burst frequency, burst duration, action potentials (AP) per burst, average AP frequency, and coefficient of variation for all ISIs measured. n = 10 wildtype and 11 *P2ry1* KO SGNs (two-tailed Student’s t-test with Bonferroni correction; ***p<0.0005, **p<0.005, ns, not significant). **(I)** Average log-binned interspike interval histograms from wildtype and *P2ry1* KO SGNs.

### P2RY1 promotes auditory neuron firing *in vivo*

The highly synchronized electrical activity exhibited by IHCs prior to hearing onset propagates through the entire developing auditory system to induce correlated firing of auditory neurons within isofrequency zones (***Babola et al., 2018***; ***Tritsch et al., 2010a***). To determine if P2RY1 is required to produce this form of correlated activity, we used *in vivo* wide-field epifluorescence microscopy of the inferior colliculus (IC) in mice that express GCaMP6s in all neurons (*Snap25-T2A-GCaMP6s* and *Snap25-T2A-GCaMP6s;P2ry1*^*–/–*^ mice). Time lapse imaging revealed that both control and *P2ry1* KO mice exhibited correlated neuronal activity confined to stationary bands oriented along the tonotopic axis (***Figure 7***A-C). Spontaneous events were less frequent in *P2ry1* KO mice (9.7 ± 0.8 events per minute compared to 13.4 ± 0.7 events per minute in control; two-tailed Student’s t test, p = 0.002), although the events were similar in amplitude and duration (half-width) (***Figure 7***D), suggesting that some compensatory amplification of events occurs in the CNS of these mice, similar to that seen in *Vglut3* KO mice (***Babola et al., 2018***). Spontaneous activity in *P2ry1* KO mice differed from controls in three other ways. First, the contralateral bias exhibited for each event was higher, with the weaker relative to stronger side amplitude decreasing from 0.61 ± 0.02 to 0.44 ± 0.02 (two-tailed Student’s t test, p = 3.0e-6) (***Figure 7***D). Second, the coefficient of variation (ratio of standard deviation to the mean) of event amplitudes was 40% higher relative to controls (***Figure 7***D). Third, a detailed examination of the spatial location of events across the tonotopic axis (***Figure 7***E) revealed that activity in brain areas later responsible for processing higher frequency tones (∼8 – 16 kHz) was reduced by 68% in *P2ry1* KO mice, while activity in low frequency areas was unaltered (***Figure 7***F-H). In *P2ry1* KO mice, bilateral removal of both cochleae abolished activity in the IC, demonstrating that activity in these mice originates in the periphery (***Figure 7–Figure Supplement 1***A-C).

**Figure 7.**
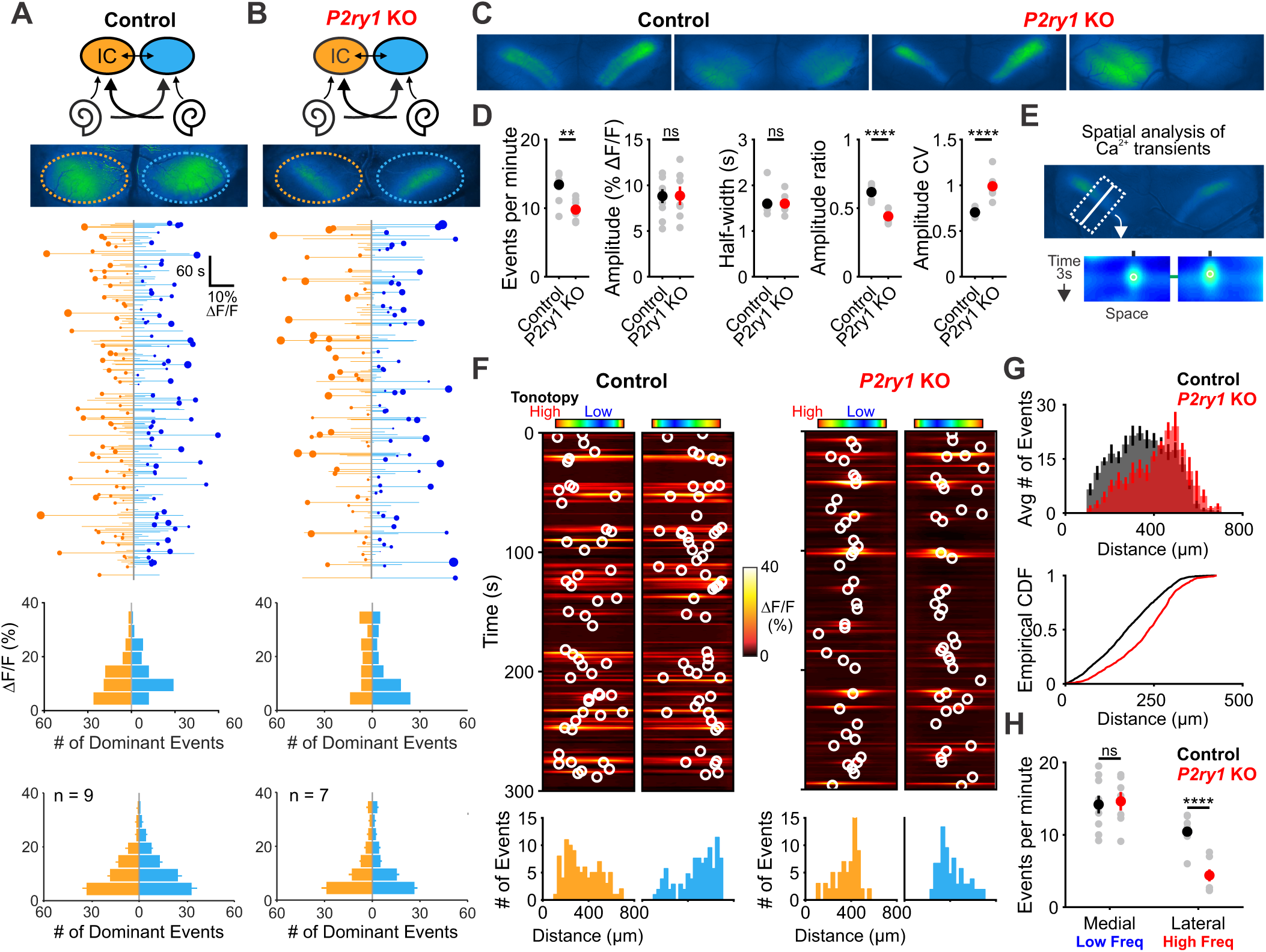
*P2ry1* KO mice exhibit reduced and spatially restricted spontaneous activity in the inferior colliculus. **(A)** Diagram illustrating flow of information through the auditory system and average intensity image over the 10 min imaging session. (middle) Activity over time in left and right IC in an individual where each line indicates the fluorescence intensity of each detected event; the circle indicates the dominant lobe, and the size of the circle indicated the difference in fluorescence. (bottom) Histograms showing the frequency of dominant events of a given amplitude for this experiment and for all experiments. Imaging was performed in *Snap25-T2A-GCaMP6s* mice (n = 9 mice). **(B)** Similar to (A), but in *Snap25-T2A-GCaMP6s;P2ry1*^*–/–*^(*P2ry1* KO) mice (n = 7 mice). **(C)** Images of spontaneous events in the IC of in control (*Snap25-T2A-GCaMP6s*) and *P2ry1* KO mice (*Snap25-T2A-GCaMP6s;P2ry1*^*–/–*^). **(D)** Comparisons of average frequency, amplitude, half-width, and event ratio from control and *P2ry1* KO mice. Bilateral amplitude ratio was calculated for events simultaneous across both lobes of the IC and defined as the ratio of the weak to the strong side amplitude. A ratio of 1 would indicate complete synchrony between lobes; a ratio of 0 would indicate complete asymmetry. n = 9 control and n = 7 *P2ry1* KO mice (two-tailed Student’s t test with Bonferroni correction; ****p<5e-4, **p<0.005, ns: not significant). **(E)** Exemplars of a single-banded event. Rectangular ROIs were placed as shown and averaged to create a ‘line-scan’ across the tonotopic axis. (bottom) Heat maps of activity as a function of time and distance; circles indicate detected peaks. **(F)** Activity over a five-minute time frame in the left and right IC of control and *P2ry1* KO mice. Circles indicate detected peaks. (bottom) Histograms of peak locations. **G)** Histogram of average number of events across all control (black) and *P2ry1* KO (red) mice. (bottom) Cumulative distribution function of event locations across the tonotopic axis pooled from all animals. Events from left and right IC were combined for each experiment. **(H)** Quantification of event frequency in the medial (low frequency) and lateral (high frequency) regions of the IC. n = 9 control and 7 *P2ry1* KO mice (two-tailed Student’s t test with Bonferroni correction; ****p<5e-5, ns, not significant).

Although *P2ry1* KO mice mimic some aspects of acute P2RY1 inhibition, the absence of P2RY1 signaling throughout life may have led to compensatory changes, such as the increase in non-purinergic ISC activity (see ***Figure 3–Figure Supplement 2***E). Therefore, to better assess the role of P2RY1 in generating spontaneous activity *in vivo*, we delivered a solution containing MRS2500 into the intraperitoneal cavity of mice while imaging activity in the IC. Compared to mice injected with control solution (5% mannitol), mice injected with MRS2500 exhibited dramatic reductions in IC event frequency (from 13.3 ± 0.8 to 3.9 ± 1.1 events per minute; two-tailed Student’s t test, p = 0.0001) and amplitude (from 9.9 ± 0.5 to 4.9 ± 0.8% ΔF/Fo; two-tailed Student’s t test, p = 0.0006) ∼5 minutes after administration (***Figure 8***A-D). This decrease was specific to the IC, as SC retinal wave activity (***Ackman et al., 2012***) was unaffected by acute MRS2500 administration (***Figure 8***B,C,E), indicating that the locus of action is likely within the cochlea, which has been shown to have a less intact blood-tissue barrier at this age (***Suzuki et al., 1998***). Spatial analysis revealed that unlike the selective deficit observed in higher frequency zones in *P2ry1* KO mice, the inhibition was not limited to certain tonotopic regions, but rather occurred evenly across all frequency zones (***Figure 8***F,G). Together, these data indicate that ISC P2RY1 autoreceptors within the cochlea play a critical role in initiating spontaneous bursts of neural activity in auditory centers within the brain prior to hearing onset.

**Figure 8.**
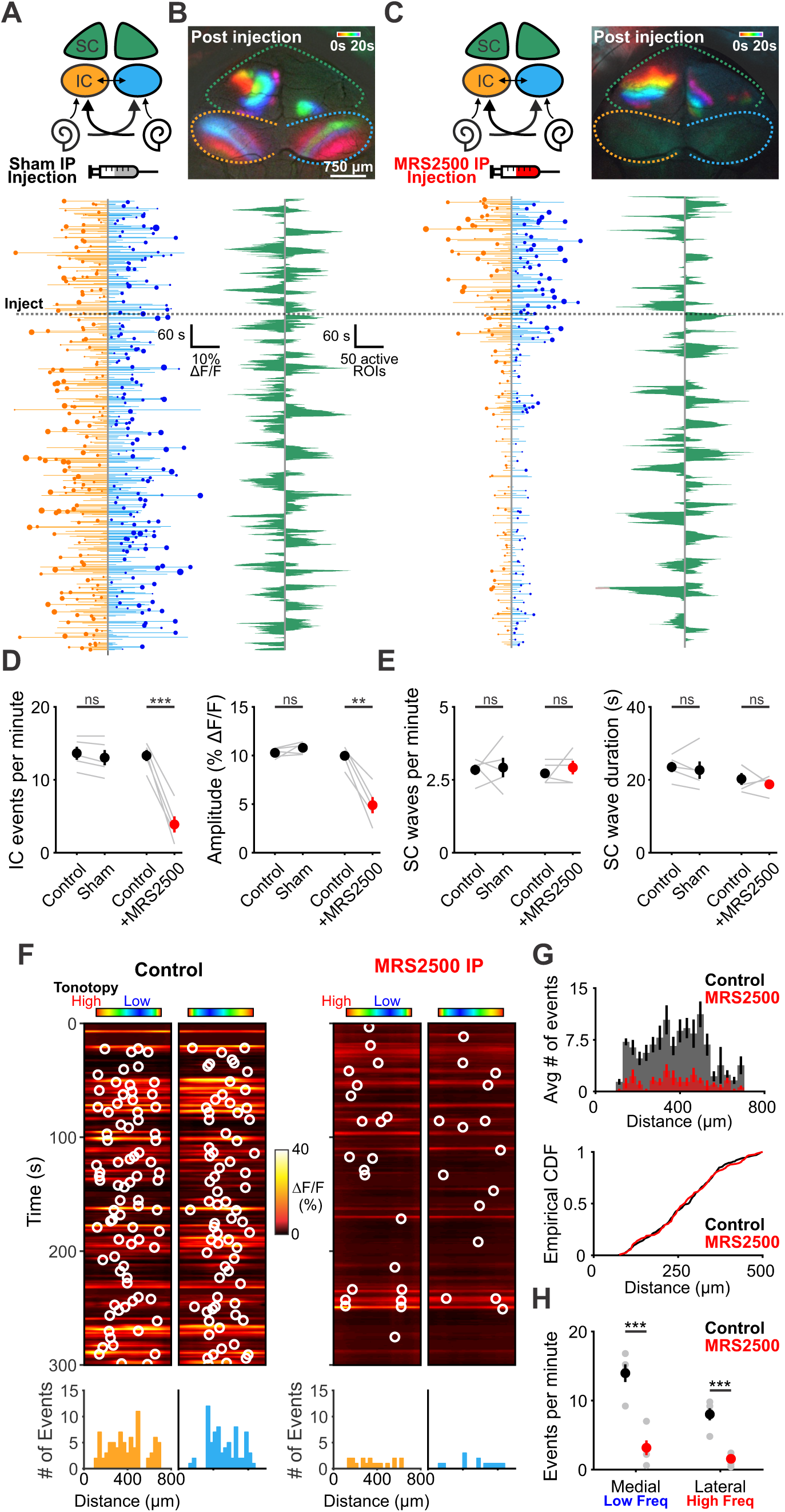
Delivery of MRS2500 *in vivo* dramatically reduces spontaneous activity in the developing auditory system. **(A)** Diagram illustrating flow of information to the midbrain and the visual superior colliculus. Sham solution (5% mannitol) was injected via IP catheter during imaging. (bottom) Activity over time in left and right IC in an individual. Each line indicates a detected event, circle indicates the dominant lobe, and the size of the circle indicates the difference in fluorescence. Dashed line is time of injection. Caption continued on next page. **(B)** Calcium transients in the midbrain, color-coded based on time of occurrence following sham injection. (bottom) Calcium transients observed in the left and right SC. **(C)** Similar to (A) and (B), but with injection of MRS2500 (50µL of 500µM MRS2500 in 5% mannitol solution). **(D)** Plot of IC event frequency and amplitude in sham and MRS2500 injected animals. n = 5 mice for each condition (two-tailed paired Student’s t test with Bonferroni correction; ***p<0.005, **p<0.005, ns: not significant). **(E)** Plot of SC wave frequency and duration in sham and MRS2500 injected animals. n = 5 mice for each condition (two-tailed paired Student’s t test with Bonferroni correction; ns: not significant). **(F)** Activity along the tonotopic axis over a five-minute time frame in the left and right IC before (left) and after (right) MRS2500 injection. Circles indicate detected peaks. (bottom) Histograms of peak locations. **(G)** Histogram of average number of events before (black) and after (red) MRS2500 injection. (bottom) Cumulative distribution function of event locations across the tonotopic axis pooled from all animals. Events from left and right IC were combined for each experiment. **(H)** Quantification of event frequency in the medial (low frequency) and lateral (high frequency) regions of the IC. n = 5 mice (two-tailed Student’s t test with Bonferroni correction; ***p<0.005).

## Discussion

Intense periods of neuronal activity dramatically alter the ionic composition of the extracellular environment, leaving behind excess K^+^ that can alter neuronal excitability, induce spontaneous activity and trigger debilitating seizures. In the CNS, homeostatic control of extracellular K^+^ levels is accomplished by glial cells, which redistribute K^+^ passively through ion channels and actively through facilitated transport, but much less is known about the mechanisms that control excitability in the peripheral nervous system. Sensory hair cells and primary auditory neurons in the cochlea are surrounded by supporting cells that share key features with CNS glia and are thought to redistribute K^+^ that accumulates during sound detection. However, prior to hearing onset, ATP dependent K^+^ release from these cells triggers periodic bursts of activity in nearby IHCs that propagate throughout the auditory system. Here, we demonstrate that this form of intrinsically generated activity is initiated through activation of P2RY1, a G_q_-coupled metabotropic purinergic receptor. Acute inhibition or genetic removal of this receptor dramatically reduced spontaneous activity and disrupted burst firing in IHCs, SGNs and central auditory neurons. In addition to triggering episodic K^+^ dependent depolarization of hair cells, activation of P2RY1 also enhanced K^+^ clearance by increasing the volume of extracellular space, allowing more rapid dissipation of extracellular K^+^ transients. This duality of purpose, to induce K^+^ efflux and enhance K^+^ clearance, promotes discrete bursts of activity throughout the developing auditory system.

### Purinergic signaling in the developing cochlea

Before the onset of hearing, neurons in the auditory system that will process similar sound frequencies exhibit periodic bursts of highly correlated activity, an entrainment that is initiated within the cochlea (***Babola et al., 2018***; ***Clause et al., 2014***; ***Sonntag et al., 2009***; ***Tritsch et al., 2010a***). Within the developing cochlear epithelium, spontaneous release of ATP from ISCs activates purinergic receptors, triggering a rapid increase of intracellular Ca^2+^, gating of TMEM16A Ca^2+^-activated Cl^−^ channels, and subsequent Cl^−^ and K^+^ efflux into the extracellular space (***Tritsch et al., 2007***; ***Wang et al., 2015***). This transient K^+^ efflux is sufficient to depolarize nearby IHCs, resulting in a burst of Ca^2+^ action potentials, release of glutamate, and suprathreshold activation of postsynaptic SGNs via AMPA and NMDA receptors (***Tritsch et al., 2010a***; ***Zhang-Hooks et al., 2016***). Our results show that activation of metabotropic P2RY1 autoreceptors is a key first step in this transduction pathway. P2RY1 is highly expressed by ISCs at a time when spontaneous activity is prominent in the cochlea (***Scheffer et al., 2015***; ***Tritsch and Bergles, 2010***) (***Figure 2***A), and spontaneous activity was reduced when intracellular Ca^2+^ stores were depleted or PLC was inhibited (***Figure 1***B-F), manipulations that disrupt canonical G_q_-coupled GPCR signaling pathways (***Erb and Weisman, 2012***; ***Fabre et al., 1999***). Moreover, our pharmacological studies indicate that P2RY1 is both necessary and sufficient for spontaneous current generation in supporting cells (***Figure 3***B, ***Figure 4***B), and inhibition of P2RY1 *in vivo* profoundly decreased cochlea-generated activity in the auditory midbrain (***Figure 8***C). This reliance on P2RY1 is unexpected, as ionotropic P2X receptors are also widely expressed in the developing cochlea (***Brändle et al., 1999***; ***Lahne and Gale, 2008***; ***Liu et al., 2015***; ***Nikolic et al., 2003***; ***Scheffer et al., 2015***; ***Tritsch et al., 2007***). The lack of P2X engagement may reflect the particular spatial-temporal characteristics of ATP release by ISCs, which may occur in locations devoid of P2X receptors or yield ATP concentration transients that favor P2RY1 activation. Exogenous ATP can induce all of the phenomenon associated with spontaneous events (ISC currents, crenation, IHC depolarization, SGN burst firing); however, it is possible that other nucleotides are released that have greater affinity for P2RY1 (e.g. ADP), or that extracellular nucleotidases rapidly convert ATP to ADP/AMP that favor activation of these native metabotropic receptors (***von Kügelgen, 2006***; ***Vlajkovic et al., 1998, 2002***).

### Control of extracellular K^+^ dynamics by supporting cells

Pharmacological inhibition of P2RY1 unexpectedly induced IHCs to gradually depolarize and begin tonic, uncorrelated firing, a phenotype also observed in *P2ry1* KO mice (***Figure 5–Figure Supplement 1***C). Our studies indicate that this phenomenon occurs because P2RY1 controls the volume of the extracellular space in the organ of Corti. Activation of P2RY1 induces ISCs to shrink osmotically (crenate), a consequence of ion and water efflux that is triggered by opening of TMEM16A channels (***Figure 4***G). The resulting increase in extracellular space lasts for many seconds and enhances dissipation of extracellular K^+^, visible through the time-dependent shift in the reversal potential of K^+^ mediated tail currents (***Figure 4***G,I). Conversely, inhibition of P2RY1 increased the size of ISCs, a swelling-induced “relaxation” that concomitantly decreased extracellular space around IHCs. K^+^ accumulation and depolarization of IHCs followed, an effect absent when IHC K^+^ channels were inhibited (***Figure 3***J,K). This phenomenon is consistent with the depolarizing shift in the resting membrane potential of IHCs observed in *Tmem16A* cKO mice (***Wang et al., 2015***), which similarly blocks ISC crenation, and with studies in the brain where inducing cell swelling with hypoosmotic solutions or impairing K^+^ buffering results in neuronal epileptiform activity (***Larson et al., 2018***; ***Murphy et al., 2017***; ***Thrane et al., 2013***). Basal P2RY1 activation in supporting cells therefore hyperpolarizes nearby IHCs in the developing cochlea by expanding the extracellular space and lowering local K^+^ concentrations. These changes increase the dynamic range of IHCs and allow finer control of excitability through transient ATP mediated signaling events.

The tonic inward current that develops in ISCs in response to P2RY1 block was abolished when homeostatic K^+^ release pathways (Na^+^ channels, Ca^2+^ channels, Na^+^-K^+^-Cl^−^cotransporters, and Na,K-ATPase) were inhibited (***Figure 3***H,I), suggesting that K^+^ redistribution mechanisms, in the absence of ISC crenation, are weak at this stage of development. Indeed, although the membrane potential of ISCs is close to E_K_, their membrane conductance is dominated by intercellular gap junction channels; when uncoupled from their neighbors, they exhibit very high (1–2GΩ) input resistance (***Jagger and Forge, 2014***; ***Wang et al., 2015***), suggesting that few K^+^ leak channels are expressed. The presence of tight junctions at the apical surface of the cochlear epithelium and the limited K^+^ conductance of ISCs may restrict passive diffusion and dilution of K^+^, similar to what has been described in the vestibular epithelium (***Contini et al., 2017***), thus necessitating uptake via alternative mechanisms. Both inner phalangeal and Dieters’ cells (which envelop the inner and outer hair cells, respectively) express K^+^-Cl^−^ symporters, Na,K-ATPase pumps, and inwardly-rectifying K^+^ channels that may siphon K^+^ into the supporting cell syncytium after extrusion from hair cells. However, the apparently low capacity of these systems places a greater dependence on diffusion within the extracellular volume fraction controlled by the supporting cells.

In the CNS, astrocytes facilitate rapid dissipation of accumulated K^+^ through the glial syncytium via gap junctions (***Kofuji and Newman, 2004***), a mechanism termed spatial buffering. Astrocytes are effcient K^+^ sinks due to their highly negative resting potential (∼–85 mV) and large resting K^+^ conductance dominated by inward rectifying K^+^ channels and two-pore K^+^ channels (***Ryoo and Park, 2016***; ***Olsen, 2012***). While uptake of K^+^ through these conductances is passive, recent studies suggest that K^+^ buffering in astrocytes is also actively regulated by purinergic receptors. Following stimulation of native astrocyte purinergic receptors or foreign G_q_-coupled receptors (MrgA1) and release of Ca^2+^ from intracellular stores, Na,K-ATPase activity increased, resulting in a transient decrease in extracellular K^+^, hyperpolarization of nearby neurons, and reduction in their spontaneous activity (***Wang et al., 2012***). Although P2RY1 is expressed by some astrocytes and can trigger Ca^2+^ waves (***Gallagher and Salter, 2003***), this mechanism does not appear to regulate IHC excitability in the cochlea, as stimulation of P2RY1 in *Tmem16a* cKO mice, which have intact metabotropic receptor signaling but no crenations (***Wang et al., 2015***), did not hyperpolarize IHCs (***Figure 4***I,J). Thus, astrocytes and cochlear ISCs use purinergic signaling in different ways to maintain the ionic stability of the extracellular environment and control the excitability of nearby cells.

### Role of supporting cells in the generation of spontaneous activity

Our understanding of how non-sensory cells contribute to spontaneous activity has been limited by a lack of *in vivo* mechanistic studies. Recent advances in visualizing cochlea-induced spontaneous activity in central auditory centers *in vivo* using genetically-encoded calcium indicators (***Babola et al., 2018***) allowed us to assess whether supporting cell purinergic receptors are involved in generating this activity. Prior to hearing onset, the blood-labyrinth barrier within the inner ear is not fully formed (***Suzuki et al., 1998***), permitting pharmacological access to the cochlea at this age. Infusion of a P2RY1 inhibitor into the intraperitoneal space dramatically decreased activity within the inferior colliculus within minutes, while retina-induced activity in the superior colliculus (***Ackman et al., 2012***) was unaffected (***Figure 8***), suggesting that inhibition is not due to activation of astrocyte P2RY1 receptors; as noted above, inhibition of P2RY1 in astrocytes would be expected to enhance, rather than inhibit neuronal activity (***Wang et al., 2012***).

*In vivo* imaging in *P2ry1* KO mice recapitulated many aspects of changes seen when P2RY1 was acutely inhibited, with significantly reduced neuronal activity observed in lateral regions of the IC (later active to 8-16kHz tones; ***Figure 7***). However, neuronal burst firing persisted within central regions of the IC, regions that will ultimately process lower frequency sounds (3-8kHz). Developing sensory systems exhibit a remarkable ability to preserve spontaneous activity. In the visual system, cholinergic antagonists injected directly into the eye blocks retinal waves *in vivo* (***Ackman et al., 2012***), but genetic removal of the β2 acetylcholine receptor subunit alters, but does not abolish, peripherally-generated activity (***Zhang et al., 2012***). In the auditory system, *in vivo* spontaneous activity can be blocked by acute inhibition of cochlear AMPARs, but deaf mice that lack the ability to excite SGNs (*Vglut3* KO mice) exhibit activity patterns remarkably similar to control mice (***Babola et al., 2018***). These robust homeostatic mechanisms allow spontaneous activity to persist despite disruption of key transduction components. Local purinergic signaling within the cochlea may still initiate tonotopic activity in central auditory circuits of *P2ry1* KO mice, as events in the IC exhibited spatial and temporal characteristics similar to controls. IHCs and SGNs are more depolarized in these mice, reducing the threshold for activation by other purinergic receptors. Although such gain-of-function changes in the developing nervous system present challenges for interpretation of genetic manipulations, preservation of early, patterned activity in children that carry deafness mutations may improve the outcome of later therapeutic interventions to restore hearing.

### Purinergic receptors in the adult cochlea

In the adult inner ear, members of all purinergic receptors subtypes (ionotropic P2X receptors, metabotropic P2Y, and adenosine P1 receptors) are expressed by cells throughout the sensory epithelium, Reisner’s membrane, stria vascularis, and SGNs (***Housley et al., 2009***; ***Huang et al., 2010***). The widespread expression of these receptors coupled with observations of increased endolymphatic ATP concentrations following trauma (***Muñoz et al., 1995a***) have led to the hypothesis that these receptors serve a neuroprotective role. Indeed, infusion of ATP into the inner ear profoundly reduces sound-evoked compound action potentials in the auditory nerve (***Bobbin and Thompson, 1978***; ***Muñoz et al., 1995b***), due to decreased endolymphatic potential following shunting inhibition through P2XR2 (***Housley et al., 2013***). Consistent with these observations, *P2rx2* KO mice and humans with a P2RX2 variant (c.178G > T) experience progressive sensorineural hearing loss (***Yan et al., 2013***). Ca^2+^ imaging and recordings from adult cochleae have also revealed robust responses to UTP in the inner sulcus, pillar cells, and Dieters’ cells (***Sirko et al., 2019***; ***Zhu and Zhao, 2010***), suggesting that metabotropic purinergic receptors continue to be expressed. Following traumatic noise damage, ATP release could activate K^+^ buffering mechanisms in supporting cells, enhance K^+^ redistribution, reduce IHC depolarization and prevent excitotoxic damage. Purinergic receptors may also contribute to IHC gain control by influencing their membrane potential, as ATP circulates in the endolypmph at low nanomolar concentrations (***Muñoz et al., 1995a***). Further studies involving conditional deletion of *P2ry1* from ISCs in the adult cochlea may help to define the role of this receptor in both normal hearing and injury contexts.

## Methods and Materials

Both male and female mice and rats of postnatal days P6–P15 were used for all experiments and randomly allocated to experimental groups. Transgenic breeders were crossed to female FVB mice to improve litter survival. Mice were housed on a 12 hour light/dark cycle and were provided food ad libitum. This study was performed in accordance with the recommendations provided in the Guide for the Care and Use of Laboratory Animals of the National Institutes of Health. All experiments and procedures were approved by the Johns Hopkins Institutional Care and Use Committee (protocol M018M350). All surgery was performed under isoflurane anesthesia and every effort was made to minimize suffering.

### Electrophysiology

For inner supporting cell recordings, apical segments of the cochlea were acutely isolated from P6-P8 rat (***Figure 1***) and mouse pups (all other figures) and used within 2 hours of the dissection. Cochleae were moved into a recording chamber and continuously superfused with bicarbonate-buffered artificial cerebrospinal fluid (1.5–2mL/min) consisting of the following (in mM): 119 NaCl, 2.5 KCl, 1.3 MgCl_2_, 1.3 CaCl_2_, 1 NaH_2_PO_4_, 26.2 NaHCO_3_, 11 D-glucose and saturated with 95% O_2_ / 5% CO_2_ to maintain a pH of 7.4. Solutions were superfused at either room temperature or near physiological temperature (32–34°C) using a feedback-controlled in-line heater (Warner Instruments), as indicated in figure legends. Whole-cell recordings of inner supporting cells (ISCs) were made under visual control using differential interference contrast microscopy (DIC). Electrodes had tip resistances between 3.5–4.5MΩ when filled with internal consisting of (in mM): 134 KCH_3_SO_3_, 20 HEPES, 10 EGTA, 1 MgCl_2_, 0.2 Na-GTP, pH 7.3. Spontaneous currents were recorded with ISCs held at –80mV.

For inner hair cell recordings, apical segments of the cochlea were acutely isolated from P6-P8 mouse pups and used within 2 hours of the dissection. Cochleae were moved into a recording chamber and continuously superfused with bicarbonate-buffered artificial cerebrospinal fluid (1.5– 2mL/min) consisting of the following (in mM): 115 NaCl, 6 KCl, 1.3 MgCl_2_, 1.3 CaCl_2_, 1 NaH_2_PO_4_, 26.2 NaHCO_3_, 11 D-glucose. Solutions were saturated with 95% O_2_ / 5% CO_2_ to maintain a pH of 7.4. Solutions were superfused at room temperature. Electrodes had tip resistances between 4.5– 6.0MΩ when filled with internal consisting of (in mM): 134 KCH_3_SO_3_, 20 HEPES, 10 EGTA, 1 MgCl_2_, 0.2 Na-GTP, pH 7.3. For hair cell recordings with K^+^ channels inhibited with cesium and TEA, the internal solution consisted of (in mM): 100 cesium methanesulfonate, 20 TEA-Cl, 10 EGTA in CsOH, 20 HEPES, 1 MgCl_2_, 0.2 Na-GTP, pH 7.3 with CsOH. Spontaneous currents were recorded with IHCs held at near their resting membrane potential (–75 to –80mV).

Errors due to the voltage drop across the series resistance and the liquid junction potential were left uncompensated for recordings of spontaneous activity. For IHC recordings with K^+^ accumulation voltage protocols (Figure 4), the amplifier compensation circuit was used to compensate 70% of the access resistance. Recordings that displayed more than a 10% increase in access resistance or access resistances > 30 MΩ were discarded. ISC and IHC spontaneous currents were recorded with pClamp 10 software using a Multiclamp 700B amplifier, low pass filtered at 2kHz, and digitized at 5kHz with a Digidata 1322A analog-to-digital converter (Axon Instruments).

Action potentials were analyzed offline using custom routines written in Matlab 2017b (Mathworks). Briefly, raw traces were high-pass filtered to remove baseline drift and spikes were identified using an amplitude threshold criterion. As described previously (***Tritsch et al., 2010a***), bursts were identified by classifying interspike intervals into non-bursting intervals (> 1s), burst intervals (30ms–1s), and mini-burst intervals (< 30ms). Bursts were defined as clusters of at least 10 consecutive burst intervals (with mini-burst intervals being ignored in the context of burst detection). Spikes within mini-bursts were included when calculating the number of spikes within a burst. Colored raster plots were generated by grouping spikes into one-second bins and applying a color map to the resulting data (modified ‘hot’ colormap; Matlab).

### Cochlear explant culture

Cochleae were dissected from postnatal day 5-6 control (*P2ry1*^*+/+*^ or *Pax2-Cre;R26-lsl-GCaMP3*) and *P2ry1* KO (P2ry1–/– or *Pax2-Cre;R26-lsl-GCaMP3;P2ry1*^*–/–*^) mice in ice-cold, sterile HEPES-buffered artificial cerebrospinal fluid (aCSF) consisting of the following (in mM): 130 NaCl, 2.5 KCl, 10 HEPES, 1 NaH_2_PO_4_, 1.3 MgCl_2_, 2.5 CaCl_2_, and 11 D-Glucose. Explants were mounted onto Cell-Tak (Corning) treated coverslips and incubated at 37°C for 24 hours in Dulbecco’s modified Eagle’s medium (F-12/DMEM; Invitrogen) supplemented with 1% fetal bovine serum (FBS) and 10U/mL penicillin (Sigma) prior to recording or imaging.

### Transmitted light imaging

Cochlear segments were imaged with a Olympus 40x water immersion objective (LUMPlanFl/IR) and recorded using MATLAB and a USB capture card (EZ Cap). Difference movies were generated by subtracting frames at time t_n_ and t_n+5_ seconds using ImageJ software to generate an index of transmittance change over time. To quantify transmittance changes, a threshold of three standard deviations above the mean was applied to the values. To calculate the frequency of these events, the whole field was taken as an ROI and peaks were detected using MATLAB (findpeaks function). To calculate area of these events, a Gaussian filter (σ= 2.0) was applied to the image after thresholding and the borders detected using MATLAB (bwlabel function). The area was then calculated as the number of pixels within the border multiplied by the area scaling factor (µm/pixel)^2^ measured with a stage micrometer.

### Immunohistochemistry and X-gal Reaction

Mice were deeply anesthetized with isoflurane and perfused with freshly prepared paraformaldehyde (4%) in 0.1 M phosphate buffer. Cochleae were post-fixed for 45 minutes at room temperature and stored at 4°C until processing. For X-gal reactions, P6-P8 cochleae were removed from the temporal bone and washed 3 × 5 minutes with PBS. Tissue was then incubated for 24 hours in the dark at 37°C in X-gal working solution consisting of (in mM): 5 K^+^ ferricyanide crystalline, 5 K^+^ ferricyanide trihydrate, 2 magnesium chloride, and 0.1% X-gal (GoldBio) dissolved in DMSO. After washing 3 × 5 minutes with PBS, images of cochleae were acquired on a dissecting microscope (Zeiss Stemi 305). For immunohistochemistry, fixed tissue was washed 3 × 5 minutes in PBS, placed in 30% sucrose solution overnight, and incubated in OCT mounting medium overnight at 4°C. Ten micron thick cross-sections of the cochlea were made on a cryostat and mounted on Superfrost Plus slides (Fisher), which were then allowed to dry for 1 hour before processing. Cross-sections were incubated overnight with primary antibodies against β-gal (anti-Chicken; 1:4000, Aves) and Myosin-VIIa (anti-Rabbit; 1:500, Proteus BioSciences) for detection of β-gal and Myosin-VIIa only for qualitative analysis of the *Tecta-Cre;TdT* reporter mouseline (***Figure 4–Figure Supplement 1***). Sections were then rinsed three times with PBS and incubated for two hours at room temperature with secondary antibodies raised in donkey (Alexa-488 and Alexa-546; 1:2000, Life Technologies). Slides were washed three times in PBS (second wash with PBS + 1:10,000 DAPI), allowed to dry, and sealed using Aqua Polymount (Polysciences, Inc.). Images were captured using a laser scanning confocal microscope (LSM 510 or 880, Zeiss).

### Confocal imaging of explants

After one day *in vitro*, cochleae were moved into a recording chamber and continuously superfused with bicarbonate-buffered artificial cerebrospinal fluid (1.5 - 2mL/min) consisting of the following (in mM): 119 NaCl, 2.5 KCl, 1.3 MgCl_2_, 1.3 CaCl_2_, 1 NaH_2_PO_4_, 26.2 NaHCO_3_, 11 D-glucose, and saturated with 95% O_2_ / 5% CO_2_ to maintain a pH of 7.4. A piezo-mounted objective was used to rapidly alternate between SGN cell bodies and ISCs/IHCs. Images were captured at 1 frame per second using a Zeiss laser scanning confocal microscope (LSM 710, Zeiss) through a 20X objective (Plan APOCHROMAT 20x/1.0 NA) at 512 × 512 pixel (354 × 354µm; 16-bit depth) resolution. Sections were illuminated with a 488nm laser (maximum 25mW power). MRS2500 (1µM), Tocris) was applied by addition to the superfusing ACSF.

### Analysis of *in vitro* Ca^2+^ transients

Images were imported into ImageJ and image registration (MultiStackReg) was used to correct for drifts in the imaging field. Since images were obtained at two different z-planes, images were combined into one stack for analysis. This was done by eliminating the empty bottom half of the imaging field containing ISCs and IHCs and the empty top half of the field containing SGN cell bodies and merging the two images. For analysis of coordinated activity throughout the cochlea, regions of interest were drawn around the entirety of ISCs, IHCs, and SGNs. Fluorescence changes were normalized as ΔF/F_o_ values, where ΔF = F - F_o_ and F_o_ was defined as the fifth percentile value for each pixel. Peaks in the signals were detected in MATLAB using the built-in peak detection function (findpeaks) with a fixed value threshold criterion (mean + 3 standard deviations for each cell).

To quantify frequency and areas of Ca^2+^ transients, a threshold of three standard deviations above the mean was applied to each pixel within the ROI. To calculate the frequency of these events, the whole field was taken as an ROI and peaks were detected using MATLAB (findpeaks function) on the number of thresholded pixels per frame. To calculate area of these events, a Gaussian filter (σ= 2.0) was applied to the image after thresholding and the borders detected using MATLAB (bwlabel function). The area was then calculated as the number of pixels within the border multiplied by an area scaling factor (1µm/pixel)^2^ measured with a stage micrometer.

For correlation analysis, ROIs were drawn around every IHCs in the field of view. Pairwise correlation coefficients were performed between every hair cell pair and represented as correlation matrices.

### Installation of cranial windows

Inhalation anesthesia was induced with vaporized isoflurane (4% for 5 minutes, or until mice are non-responsive to toe-pinch) and surgical plane maintained during the procedure (with 1-2% isoflurane) with a stable respiration rate of 80 breaths per minute. A midline incision beginning posterior to the ears and ending just anterior to the eyes was made. Two subsequent cuts were made to remove the dorsal surface of the scalp. A headbar was secured to the head using super glue (Krazy Glue). Fascia and neck muscles overlying the interparietal bone were resected and the area bathed in sterile, HEPES-buffered artificial cerebrospinal fluid that was replaced as necessary throughout the surgery. Using a 28G needle and microblade, the sutures circumscribing the interparietal bone were cut and removed to expose the midbrain. The dura mater was removed using fine scissors and forceps, exposing the colliculi and extensive vasculature. A 5 mm coverslip (CS-5R; Warner Instruments) was then placed over the craniotomy, the surrounding bone was dried using a Kimwipe, and super glue was placed along the outer edges of the coverslip for adhesion to the skull. Replacement 0.9% NaCl solution was injected IP and a local injection of lidocaine was given to the back of the neck. Animals were weaned off isoflurane, placed under a warming lamp, and allowed to recover for a minimum of 1 hour prior to imaging. Spontaneous activity was not seen in deeply anesthetized animals and emerged 30 minutes after recovery from isoflurane exposure, as reported previously (***Ackman et al., 2012***).

### *In vivo* calcium imaging

After 1 hour of post-surgical recovery from anesthesia, pups were moved into a swaddling 15 mL conical centrifuge tube. The top half of this tube was removed to allow access to the headbar and visualization of the midbrain. Pups were head-fixed and maintained at 37°C. using a heating pad and temperature controller (TC-1000; CWE). During the experiments, pups were generally immobile; however, occasional limb and tail twitching did occur.

For wide field epifluorescence imaging, images were captured at 10 Hz using a Hamamatsu ORCA-Flash4.0 LT digital CMOS camera attached to a Zeiss Axio Zoom.V16 stereo zoom microscope. For midbrain imaging, a 4 × 4mm field of view was illuminated continuously with a mercury lamp (Zeiss Illuminator HXP 200C) and visualized through a 1X PlanNeoFluar Z 1.0x objective at 17x zoom. Images were captured at a resolution of 512 × 512 pixels (16-bit pixel depth) after 2 × 2 binning to increase sensitivity. Each recording consisted of uninterrupted acquisition over 10 minutes or 20 minutes if injected with pharmacological agents.

### Catheterization of animals for *in vivo* imaging

After induction of anesthesia and before installing the cranial window, a catheter was placed in the intraperitoneal (IP) space of neonatal mouse pups. A 24G needle was used to puncture the peritoneum and a small-diameter catheter (SAI Infusion Technologies, MIT-01) was placed. A drop of Vetbond secured the catheter to the pup’s belly. Installation of cranial window proceeded as described above. Imaging sessions consisted of 5 minutes of baseline activity measurements, followed by a slow push of either 50µL of sham (5% mannitol solution) or MRS2500 solution (500µM in 5% mannitol solution). Imaging was continuous throughout and 20 minutes of activity total were collected. No discernable diminishment of activity was observed in sham animals.

### Image processing

For wide field imaging, raw images were imported into the ImageJ environment and corrected for photobleaching by fitting a single exponential to the fluorescence decay and subtracting this component from the signal (Bleach Correct function, exponential fit). Images were then imported into MATLAB (Mathworks) and intensities were normalized as DF/F_o_ values, where DF = F - F_o_ and F_o_ was defined as the fifth percentile value for each pixel. Ovoid regions of interest (ROIs) encompassing the entire left and right inferior colliculi were drawn. Across all conditions, the size of the ROIs was invariant, however, due to small differences in the imaging field between animals, the ROIs were placed manually for each imaging session. Peaks in the signals were detected in MATLAB using the built-in peak detection function (findpeaks) using a fixed value threshold criterion; because fluorescence values were normalized, this threshold was fixed across conditions (2% ΔF/F_o_). Occasionally, large events in the cortex or superior colliculus would result in detectable fluorescence increases in the IC. These events broadly activated the entire surface of the IC and did not exhibit the same spatially-confined characteristics as events driven by the periphery. These events were not included in the analysis.

### Analysis of spatial distribution of activity in the IC

As shown in ***Figure 7***D, a rectangle of size 125 × 50 pixels was placed perpendicular to the tonotopic axis of the IC (±55° rotation, respectively). The columns of the resulting matrix were averaged together to create a line scan (125 pixels x 1 pixel) for the entire time series. Peaks were detected using MATLAB’s imregionalmax function with a constant threshold of 3% ΔF/F_o_ across all animals. Histograms of events along the tonotopic axis were generated by summing the number of events in 25µm bins. Lateral and medial designations were assigned by splitting the area evenly between the lateral edge and the location of defined single-band events in the medial portion of the IC. Events detected on the medial edge of single-band events, reflective of the bifurcation of this information, was not included in the medial/lateral analysis.

### Analysis of retinal wave activity in the superior colliculus

ROIs (200 × 150 pixels) were placed over each lobe of the superior colliculus and downsampled by a factor of five. Signals were normalized as ΔF/F_o_ values, where ΔF = F - F_o_ and F_o_ was defined as the fifth percentile value for each pixel. In order to eliminate periodic whole-sample increases in fluorescence, the mean intensity of all pixels was subtracted from each individual pixel. Following this, pixels were considered active if they exceeded the mean + 3 standard deviations. For each point in time, the number of active pixels was summed. Retinal waves were defined as prolonged periods (> 1 second), where more than 5 pixels were active simultaneously. Retinal wave durations were defined as the total continuous amount of time that more than 5 pixels were active. Frequencies and durations are similar to earlier reports (***Ackman et al., 2012***).

### Generation of the Tecta-Cre mouseline

A crRNA (TAATGATGAATAATTCATCC) targeted near exon 2 of the Tecta gene, tracrRNA, Cas9 recombinase, and a donor plasmid containing an iCre-WPRE-polyA sequence (500 base pair homology arms) were injected into single-cell embryos that were then transferred to pseudopregnant recipient mothers. After birth, mouse pups were screened for insertion of the gene at the correct locus with two pairs of primers: one pair amplified DNA beginning 5’ of the 5’ homology arm and ending within the Cre sequence and the other amplified DNA within the polyA sequence and ending 3’ of the 3’ homology arm. These primers were then used to sequence the junctions. Of these, all mice used for experiments were derived from a single founder that was positive for both sets of primers and had 100% sequence validation. Mice were crossed to a TdTomato reporter line to examine cell-specific recombination (***Figure 4–Figure Supplement 1***).

### Quantification and statistical analysis

All statistics were performed in the MATLAB (Mathworks) programming environment. All statistical details, including the exact value of n, what n represents, and which statistical test was performed, can be found in the figure legends. To achieve statistical power of 0.8 with of a 30% effect size with means and standard deviations similar to those observed in previous studies, power calculations indicated that 7 animals in each condition were necessary (µ_1_ = 10, µ_2_ = 7, σ = 2, sampling ratio = 1). While this number was used as a guide, power calculations were not explicitly performed before each experiment; many experiments had much larger effect sizes and sample sizes were adjusted accordingly. For transparency, all individual measurements are included in the figures. Unless otherwise noted, data are presented as mean ± standard error of the mean. All datasets were tested for Gaussian normality using the D’Agostino’s K^2^ test. For single comparisons, significance was defined as p <= 0.05. When multiple comparisons were made, the Bonferroni correction was used to adjust p-values accordingly to lower the probability of type I errors. For multiple condition datasets, one-way ANOVAs were used, followed by Tukey’s multiple comparison tests.

## Acknowledgments

We thank Dr. M. Pucak and N. Ye for technical assistance, T. Shelly for machining expertise, and members of the Bergles laboratory for discussions and comments on the manuscript. TAB was supported by grants from the NIH (DC016497) and by departmental training grants (NS091018, DC000023). HCW and CJK were supported by a departmental training grant (DC000023). Funding was also provided the NIH (DC008860, NS050274), Otonomy Inc., the Brain Science Institute at Johns Hopkins University, and the Rubenstein Fund for Hearing Research to DB.

## Author contributions

Travis A. Babola, Conceptualization, Data curation, Formal analysis, Funding acquisition, Investigation, Methodology, Software, Validation, Visualization, Writing–original draft, Writing–review and editing; Calvin J. Kersbergen, Data curation, Formal analysis, Investigation, Methodology, Validation, Writing–review and editing; Han Chin Wang, Methodology, Data curation, Formal analysis, Investigation, Methodology, Writing–review and editing; Dwight E. Bergles, Conceptualization, Data curation, Funding acquisition, Methodology, Project administration, Supervision, Writing–review and editing

---

**Figure 3–Figure supplement 1.**
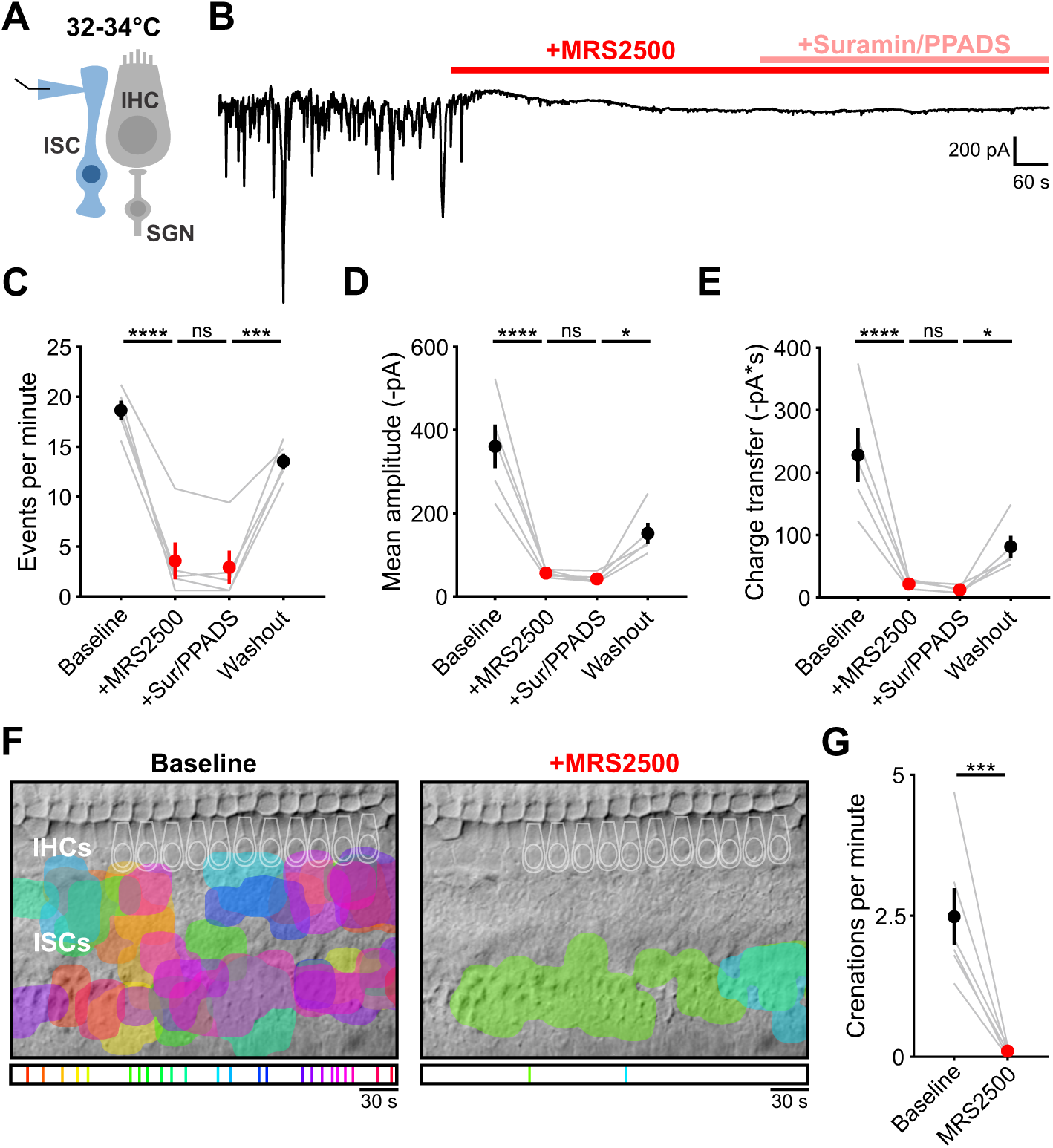
P2RY1 inhibition abolishes spontaneous inward currents near physiological temperature. **(A)** Schematic of whole-cell recording configuration from ISCs. **(B)** Spontaneous inward currents recorded from an ISC before and after application of MRS2500 (1µM) and subsequent broad spectrum purinergic antagonists suramin (10µM) and PPADS (50µM). Recordings were performed near physiological temperature (32–34°C). **(C)** Plot of event frequency. Each window measured was 5 minutes in length, washout was taken 20 minutes after superfusion of aCSF. n = 5 ISCs from 5 cochleae (one-way ANOVA; ****p<5e-5, ***p<0.0005, ns, not significant). **(D)** Plot of event amplitude. n = 5 ISCs from 5 cochleae (one-way ANOVA; ****p<5e-5, *p<0.05, ns, not significant). **(E)** Plot of average integral (charge transfer). n = 5 ISCs from 5 cochleae (one-way ANOVA; ****p<5e-5, *p<0.05, ns, not significant). **(F)** Intrinsic optical imaging performed before and after application of the P2Y1 antagonist, MRS2500 (1µM). Detected crenations are outlined in colors based on time of occurrence as indicated by timeline below image. Imaging was performed near physiological temperature (32–34°C). **(G)** Plot of crenation frequency before and after MRS2500 application. n = 6 cochleae (two-tailed paired Student’s t test; **p<0.005).

**Figure 3–Figure supplement 2.**
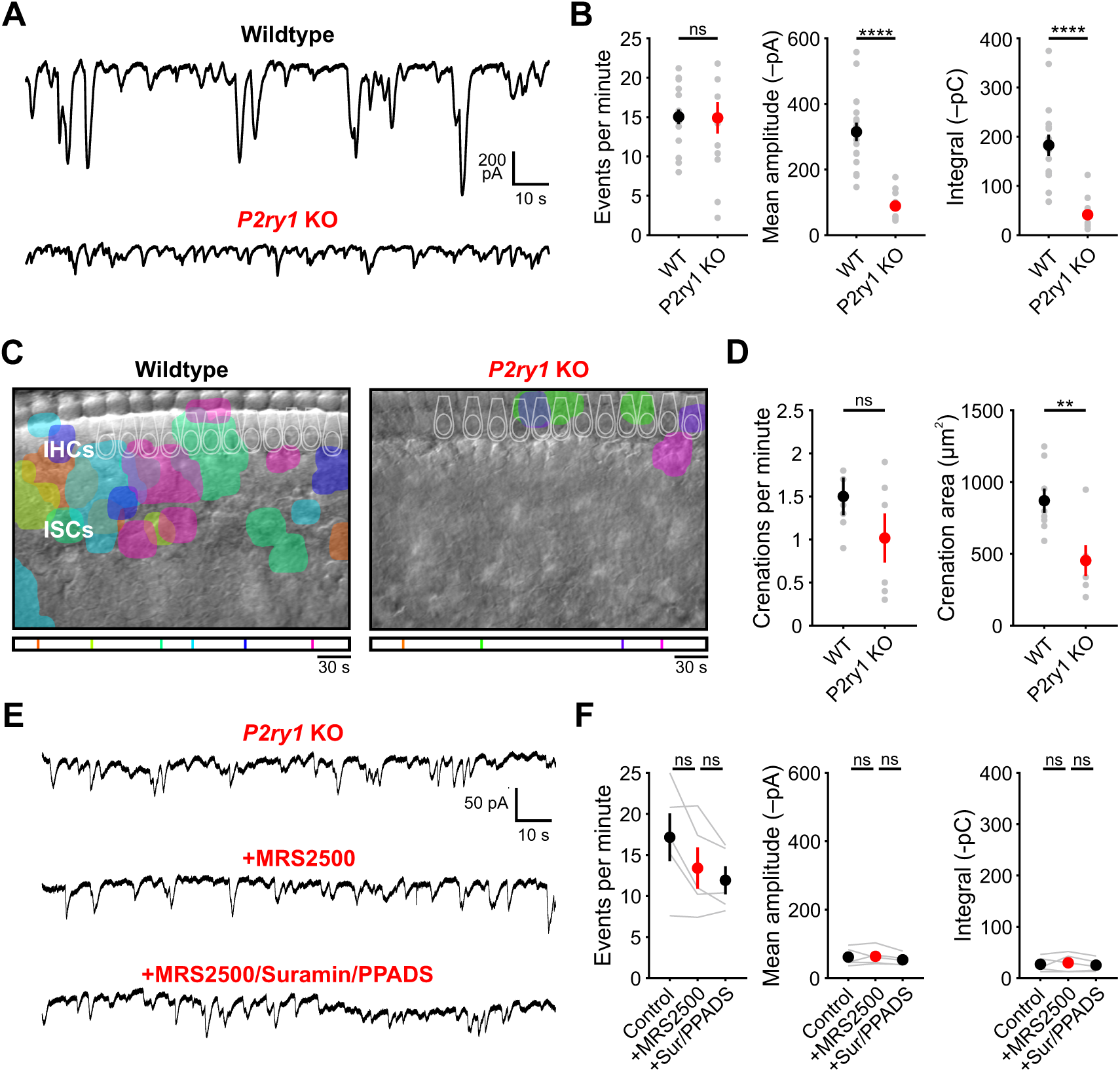
Spontaneous inward currents and crenations are dramatically reduced in *P2ry1* KO mice. **(A)** Spontaneous inward currents recorded from ISCs in wildtype and *P2ry1* KO mice. Recordings were performed near physiological temperature (32–34°C). **(B)** Plots of event frequency, amplitude, and integral (charge transfer). n = 17 wildtype and 14 *P2ry1* KO ISCs (two-tailed Student’s t-test with Bonferroni correction; ****p<0.0005, ns, not significant). **(C)** Intrinsic optical imaging performed in wildtype and *P2ry1* KO mice. Detected crenations are outlined in colors based on time of occurrence as indicated by the timeline below image. Imaging was performed at room temperature. **(D)** Plots of crenation frequency and area in wildtype and *P2ry1* KO mice. n = 8 wildtype cochleae and 6 *P2ry1* KO cochleae (two-tailed paired Student’s t test with Bonferroni correction; **p<0.005, ns, not significant). **(E)** Spontaneous inward currents recorded from an inner supporting cell in *P2ry1* KO mice before and during application of MRS2500 (1µM) and subsequent broad spectrum purinergic antagonists suramin (100µM) and PPADS (50µM). **(F)** Plots of event frequency, amplitude, and charge transfer. n = 5 *P2ry1* KO ISCs (one-way ANOVA; ns, not significant).

**Figure 4–Figure supplement 1.**
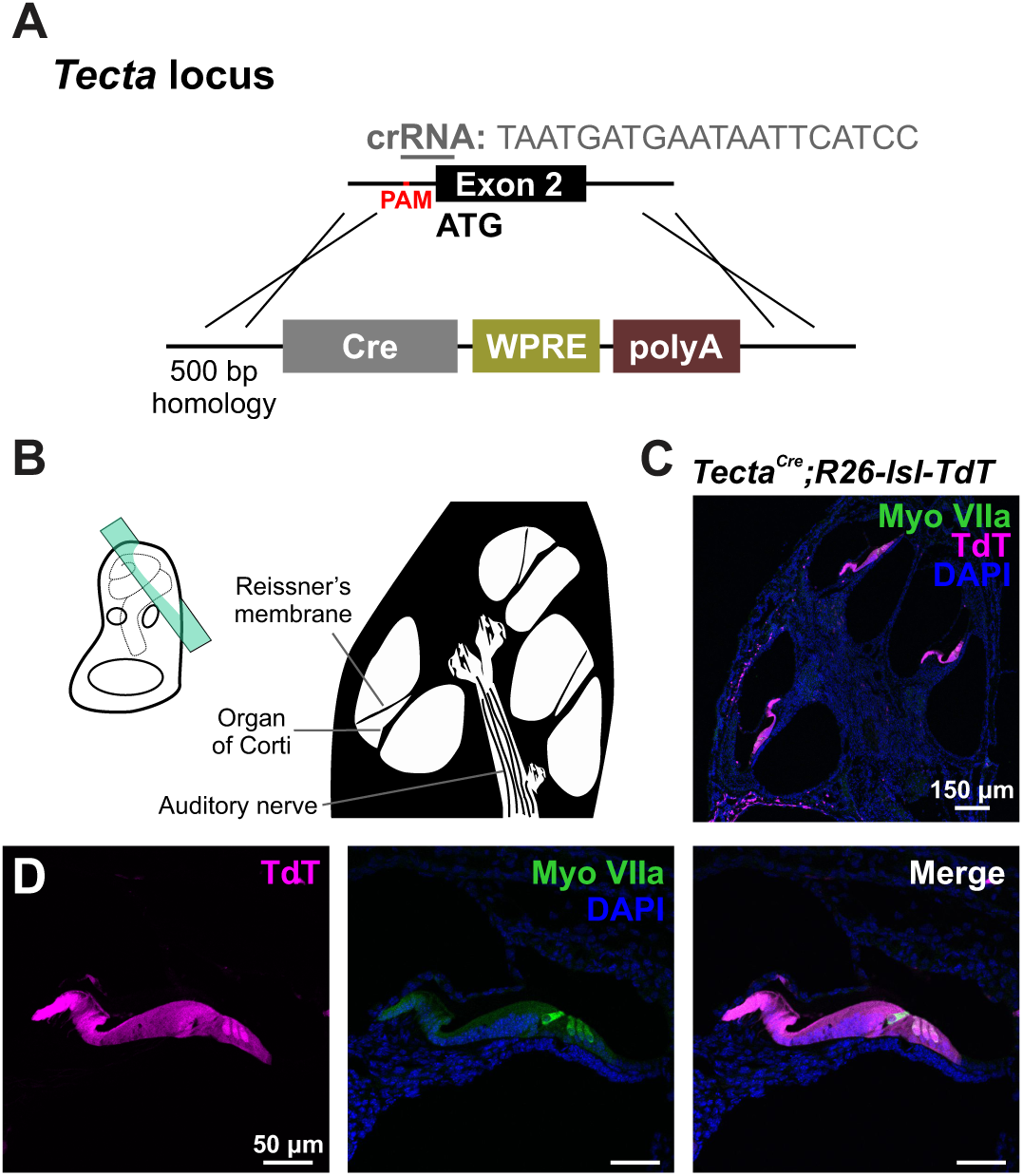
Crispr-Cas9 mediated generation of the *Tecta-Cre* mouse line. **(A)** Targeting strategy for introducing an *iCre* coding sequence into the endogenous *Tecta* locus. Note: start ATG is located in exon 2. **(B)** Schematic of temporal bone with sectioning orientation indicated with green plane. **(C)** TdTomato reporter expression observed along the entire length of a P7 cochlea. Expression was absent in stria vascularis and very sparse in apical SGNs. **(D)** TdTomato reporter expression observed in nearly all cells within the sensory epithelium, including hair cells (MyoVIIa, green).

**Figure 5–Figure supplement 1.**
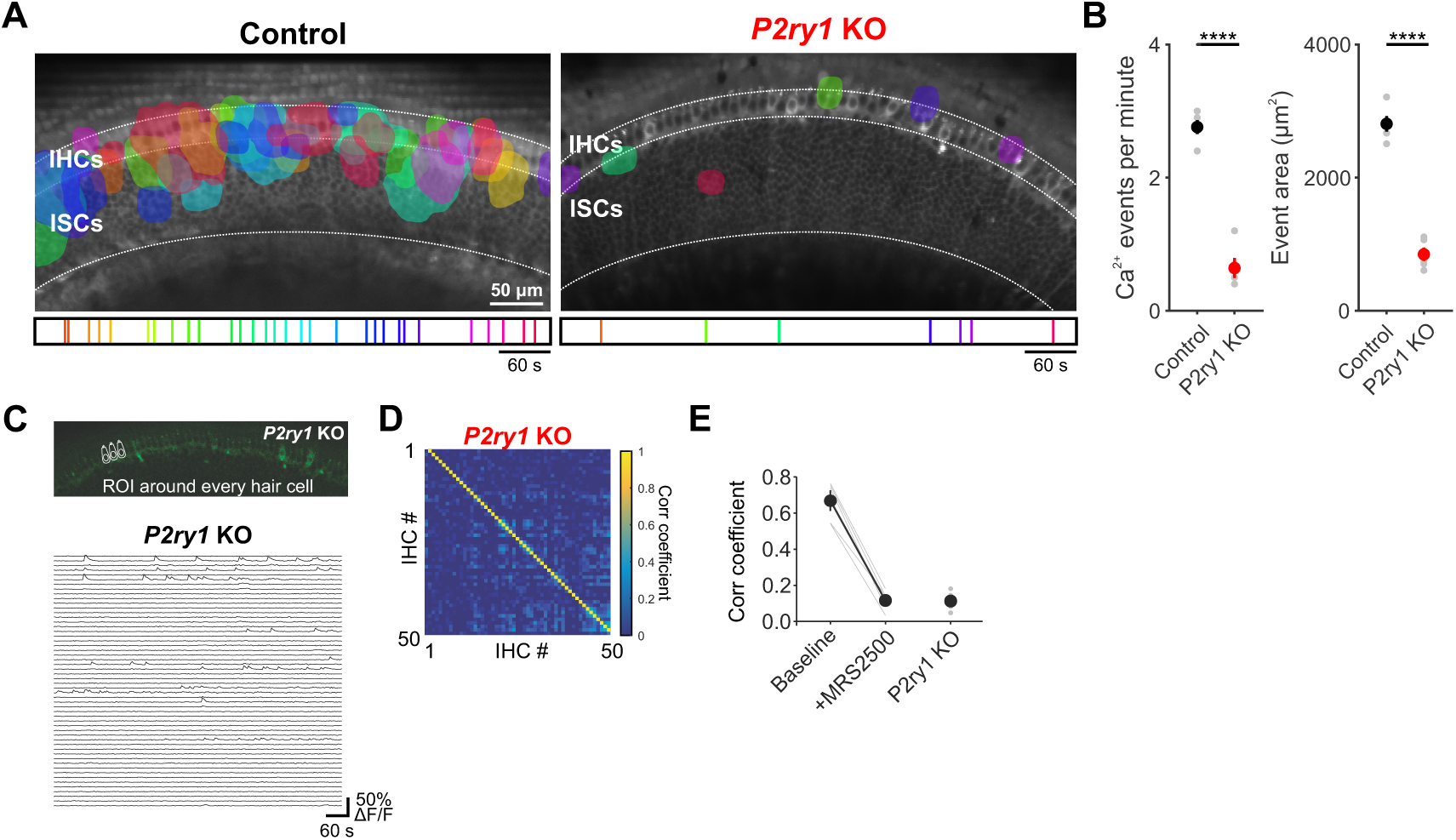
*P2ry1* KO mice exhibit reduced Ca^2+^ transients in ISCs. **(A)** Maps showing maximum area of spontaneous Ca^2+^ transients in control (*Pax2-Cre;R26-lsl-GCaMP3*) and *P2ry1* KO (Pax2-Cre;R26-lsl-GCaMP3; P2ry1–/–) mice. Ca^2+^ transients in the ISC and IHC regions are color-coded based on time of occurrence as indicated in timeline below image. Imaging was performed at room temperature. **(B)** Plots of Ca^2+^ event frequency and area in control and *P2ry1* KO mice. n = 5 control and 5 *P2ry1* KO mice (two-tailed Student’s t test with Bonferroni correction, ****p<0.05). **(C)** Exemplar images of IHC Ca^2+^ transients. ROIs were drawn around every IHC for subsequent analysis (bottom). **(D)** Correlation matrices generated by calculating the linear correlation coefficient for all IHC pairs in *P2ry1* KO mice. **(E)** Plot of average correlation coefficient among the four nearest neighboring hair cells. Data from MRS2500 experiment (**Figure 4**) is reproduced here for comparison. n = 5 cochleae (two-tailed paired Student’s t test with Bonferroni correction;***p<0.0005).

**Figure 7–Figure supplement 1.**
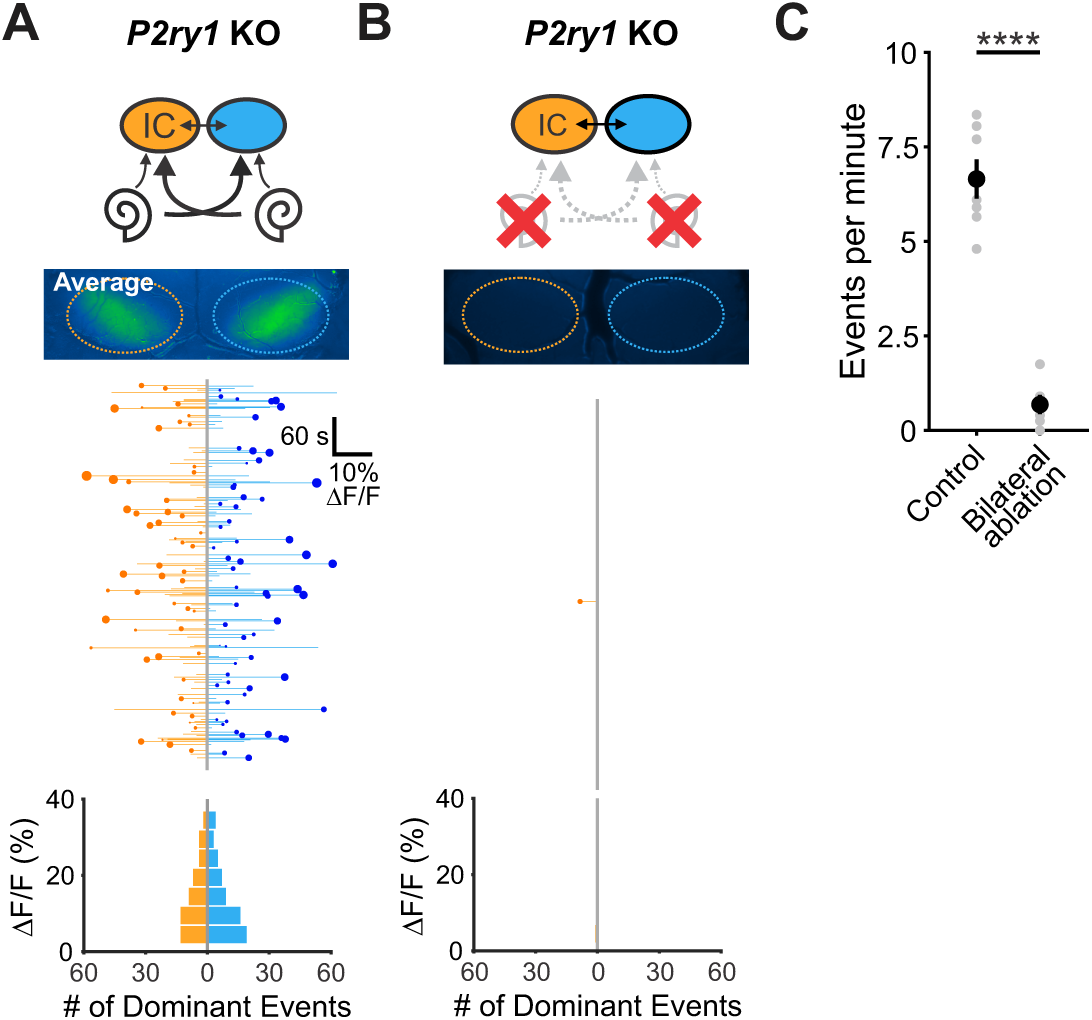
Spontaneous activity in *P2ry1* KO mice originates in the cochlea. **(A)** Top: Diagram illustrating flow of information through the auditory system and average intensity image over the 10 min imaging session. Middle: Activity over time in left and right IC in an individual where each line indicates the fluorescence intensity of each detected event; the circle indicates the dominant lobe, and the size of the circle indicated the difference in fluorescence. Bottom: Histograms showing the frequency of dominant events of a given amplitude. **(B)** Similar to A, but with bilateral ablation of the cochleae. **(C)** Plot of IC event frequency in control (*P2ry1* KO) and bilaterally ablated (*P2ry1* KO) mice. n = 7 control and n = 5 bilaterally ablated *P2ry1* KO mice (two-tailed Student’s t test; ****p<5e-5).

